# Genome-wide mapping reveals conserved and diverged R-loop activities in the unusual genetic landscape of the African trypanosome genome

**DOI:** 10.1101/357020

**Authors:** Emma Briggs, Graham Hamilton, Kathryn Crouch, Craig Lapsley, Richard McCulloch

## Abstract

R-loops are stable RNA-DNA hybrids that have been implicated in transcription initiation and termination, as well as in telomere homeostasis, chromatin formation, and genome replication and instability. RNA Polymerase (Pol) II transcription in the protozoan parasite *Trypanosoma brucei* is highly unusual: virtually all genes are co-transcribed from multigene transcription units, with mRNAs generated by linked trans-splicing and polyadenylation, and transcription initiation sites display no conserved promoter motifs. Here, we describe the genome-wide distribution of R-loops in wild type mammal-infective *T. brucei* and in mutants lacking RNase H1, revealing both conserved and diverged functions. Conserved localisation was found at centromeres, rRNA genes and retrotransposon-associated genes. RNA Pol II transcription initiation sites also displayed R-loops, suggesting a broadly conserved role despite the lack of promoter conservation or transcription initiation regulation. However, the most abundant sites of R-loop enrichment were within the intergenic regions of the multigene transcription units, where the hybrids coincide with sites of polyadenylation and nucleosome-depletion. Thus, instead of functioning in transcription termination, most *T. brucei* R-loops act in a novel role, promoting RNA Pol II movement or mRNA processing. Finally, we show there is little evidence for correlation between R-loop localisation and mapped sites of DNA replication initiation.

## Introduction

RNA-DNA hybrids display enhanced stability compared with double-stranded DNA or RNA due to the unusual conformation they adopt (1,2). Small RNA-DNA hybrids form during priming of DNA replication and within RNA polymerase (Pol) during transcription, whereas larger RNA-DNA hybrids, termed R-loops, can form when RNA exiting the RNA Pol may access the DNA before the duplex reanneals. These R-loops are exacerbated when elements of RNA biogenesis are impaired (3–7). However, R-loops are increasingly recognised as providing widespread roles (see below) (8–10), which are not all co-transcriptional, since R-loops may also form in *trans* when RNA moves from the site of its genesis to another, homologous location. One example of *trans* R-loop formation is seen during the prokaryotic CRISPR-cas defence system (11) and eukaryotic and bacterial homologous recombination proteins (which normally direct DNA repair) are capable of generating RNA-DNA hybrids (12–15).

R-loops can be detrimental to genome function, leading to instability and mutation (16–18), for instance by blocking replication or because of increased lability of the exposed single-stranded DNA. In addition, sites of DNA replication and transcription collision have been shown to accumulate R-loops (19,20). To counteract these detrimental effects, activities have been described in eukaryotic cells to limit R-loop formation during mRNA biogenesis (8,21). In addition, all cells encode activities to resolve R-loops once they form. Beyond a number of helicases that can unwind R-loops (8,21), RNase H enzymes play a key role in all cells in degrading the RNA within the hybrid. Most prokaryotes and eukaryotes encode two RNase H enzymes (22): one, eukaryotic RNase H1, is monomeric and appears conserved with bacterial RNase HI, while RNase H2 in eukaryotes is trimeric and therefore differs structurally from monomeric bacterial RNase HII. Distinct roles might be predicted by the presence of two RNase H enzymes throughout life, but separate R-loop-associated functions in the eukaryotic nucleus have been hard to identify, though RNase H2 is additionally able to excise ribonucleotides mis-incorporated into DNA (23), while RNase H1 acts on organelle genome R-loops in yeast, mammals and plants (24–26). Loss of either RNase H is lethal in mammals (25,27), but yeast mutants lacking both enzymes are viable and display increased nuclear R-loop abundance (6,24,28). Clashes between transcription and replication can be resolved in bacteria by RNase HII acting on the resulting R-loops (19), while the R-loops that form in the same circumstances in eukaryotes have been shown to lead to DNA damage signalling (20) and the recruitment of repair factors (29–31).

Growing evidence suggests that R-loops are not always detrimental, but instead are increasingly associated with functional roles. Translocations in the mammalian immunoglobulin locus can cause lymphoma, but at least some of these cancers may be an unfortunate by-product of the deliberate generation of R-loops to mediate immunoglobulin class switch recombination during B-cell maturation (32,33). R-loops also have a key role in initiating replication of bacterial plasmids (34), mitochondrial DNA (35) and phage genomes (36). Indeed, it has been suggested R-loops contribute to DNA replication initiation at cryptic bacterial origins (37) and to at least some eukaryotic nuclear DNA replication initiation (38,39). Beyond recombination and replication, gene expression control can be enacted by R-loops (8,10). In human cells R-loops at CpG island promoters protect against epigenetic silencing (40–42), while in *Arabidopsis* the hybrids may promote silencing of some genes (43). Similarly, in yeast and mammalian cells R-loop association with chromatin modification and non-coding RNA mediates termination of transcription (44–47). Indeed, R-loop and chromatin modification might be a widespread association (48–50), since the hybrids can be found throughout some protein-coding genes, rather than being limited to promoters and terminators (51,52). Finally, TERRA RNAs generated by the transcription of the telomere repeats at the ends of eukaryotic chromosome can form R-loops (53), which may provide a means to maintain telomeres in the absence of telomerase, including through recombination (54,55).

The above, widespread localisation of R-loops has only to date been explored in organisms that follow conventional rules for eukaryotic protein-coding gene expression, where each protein is normally found as a single transcription unit with its own promoter and terminator. Gene expression in kinetoplastids, a grouping of protozoans that includes several major human and animal parasites, does not conform to these rules (56). One such kinetoplastid is *Trypanosoma brucei*, and co-ordination of gene expression here largely reflects the broader range of organisms in this lineage. Virtually every protein-coding gene (~8000) in *T. brucei* is transcribed from a small number (~200) of multigene transcription units, with little evidence for functional grouping within the units (57). Only two genes have been described to have introns (58) and mature mRNAs are generated from multigene RNA transcripts by coupled trans-splicing and polyadenylation (59). Remarkably, some protein-coding genes in *T. brucei* are expressed from multigene units transcribed by RNA Pol I, where the promoters share some homology with those at rRNA gene clusters (60). The vast majority of protein coding genes are more conventionally transcribed by RNA Pol II, but from promoters that are still not fully understood. Sites of transcription initiation have been mapped to so-called strand switch regions (SSRs) that separate adjacent RNA Pol II transcription units (61,62), but conserved sequence motifs characteristic of eukaryotic RNA Pol II promoters have escaped detection. Instead, it appears increasingly likely that SSRs are more similar to dispersed and unregulated eukaryotic RNA Pol II promoters (63), consistent with a lack of control over transcription initiation and devolution of gene expression controls to post-transcriptional reactions, such as mRNA turnover (64). Transcription termination at the ends of multigene transcription units in *T. brucei* is even more poorly understood, though variant and modified histones have been localised to terminator SSRs, as has a modified base, called J (65,66). In fact, SSRs are not merely sites of transcription initiation, but are also the locations where DNA replication initiates, termed origins (67,68). Functional interaction between the replication and transcription machineries is suggested by RNA changes around the SSRs after RNAi against ORC1/CDC6, a subunit of the origin recognition complex, which binds all SSRs but directs DNA replication initiation (69) at only a subset of ~45 SSRs (67,68). What features distinguish origin-active SSRs from non-origin SSRs is unclear, but the wide separation of origins and their co-localisation with some SSRs may limit deleterious collisions between the DNA and RNA Pol machineries in the context of multigenic, pervasive transcription (67,70).

The highly divergent genetic landscape found in the *T. brucei* genome provides an excellent platform to ask what features of R-loop localisation and function are conserved across eukaryotes and to potentially reveal kinetoplastid or *T. brucei*-specific roles. Localisation of R-loops has been greatly aided by the generation of the monoclonal antibody S9.6, which binds RNA-DNA hybrids (71) and, to a lesser extent, double-stranded RNA (72). Here, we used S9.6 for RNA-DNA hybrid immunoprecipitation (IP) to evaluate R-loop distribution genome-wide in both wildtype bloodstream form (BSF, mammal-infective) *T. brucei* cells and in null mutants lacking RNase H1. We show that R-loops are very abundant across the *T. brucei* genome, with the most pronounced genomic localisation seen throughout the RNA Pol II multigene transcription units, where R-loops coincide most clearly with polyadenylation sites and regions of nucleosome depletion. Thus, in these locations we reveal a novel function for R-loops, which is not related to transcription termination, but to RNA processing and, potentially, to ensuring a chromatin landscape needed for the continued movement of RNA Pol. In addition, despite the divergence of *T. brucei* RNA Pol II promoters, we show significant enrichment of R-loops at sites of transcription initiation, indicating promoter-proximal RNA-DNA hybrids are not merely associated with the regulation of transcription but may be necessary for the mechanics of initiation. Beyond these novel R-loop functions, we reveal widespread conserved localisation to retrotransposon-associated sequences, rRNA and centromeres, but we find no evidence that R-loops preferentially form at DNA replication origins in *T. brucei*.

## Materials and Methods

### *T. brucei* cell lines

All cell lines used were BSF *T. b. brucei*, strain Lister 427, and were maintained in HMI-9 medium supplemented with 10% (v/v) FBS (Sigma-Aldrich, Missouri, USA) and 1% (v/v) penicillin-streptomycin solution (Gibco) at 37 °C and 5% CO2. Heterozygous (−/+) and homozygous (−/−) *Tbrh1* knockout cell lines were generated using two constructs containing cassettes of either blasticidin or neomycin resistance genes between α-β tubulin and actin intergenic regions, flanked by sequences homologous to the 5’ and 3’ UTRs of *TbRH1*, essentially as described in (Devlin et al 2016). Homologous flanking regions were PCR-amplified using the following primers: 5’ UTR CGACG*GGATCC*TTGCCTTACCCGTGTTTT and CGACG*TCTAGA*CCTTTTCTTTCCCATGGAC, 3’ UTR CGACG*CCCGGG*AGGTGTGTATGGGAATGA and CGACG*CTCGAG*GCACCACCCAGTATAGAAA.

### DRIP-seq analysis

DRIP was performed using a ChIP-IT Enzymatic Express kit (Active Motif). Briefly, ~ 2×10^8^ cells were grown to log phase before fixing in 1% formaldehyde for 5 min whilst shaking at room temperature, before 1 ml of 10X Glycine Buffer was added directly to the cells to stop fixation. Cells were then pelleted, re-suspended in Glycine Stop-Fix Solution and shaken at room temperature for 5 min. Cells were next lysed, according to the manufacturer’s protocol, allowing chromatin to be extracted and digested for 5 min with Enzymatic Shearing Cocktail at 37 °C to produce ~200 bp fragments. IP was performed overnight at 4 °C with 4.5 ng of S9.6 antibody (Kerafast).

Library preparation was performed using a TruSeq ChIP Library Preparation Kit (Illumina) and fragments of 300 bp, including adaptors, were selected with Agencourt AMPure XP (Beckman Coulter). Sequencing was performed with an Illumina NextSeq 500 platform. Reads were trimmed using TrimGalore (https://github.com/FelixKrueger/TrimGalore) under default settings before alignment to a ‘hybrid’ reference genome consisting of the TRUE-927 v5.1 core chromosome assembly, plus sequences of 14 Lister 427 VSG ES and 5 mVSG ES (73) using Bowtie2 (74) in “very-sensitive” mode. Reads with a MapQ value <1 were removed using SAMtools (75), leaving at least 30 million aligned reads per sample. The fold-change between input and DRIP read depth was determined for each sample over non-overlapping 50 bp windows using the DeepTools bamCompare tool: (library size was normalised via the SES method, foldchange was expressed as a ratio) and visualised as tracks with IGV (76). Regions with a fold-change ≥ 1.2 were considered enriched and adjacent enriched windows were combined to give the coordinates of final DRIP enriched regions.

Classification was accomplished by assessing DRIP enriched region coordinate overlap with different genomic regions: VSG subtelomeric arrays, RHS, centromeres, Pol I, Pol II and Pol III transcripts, SSRs, mVSG and VSG ES. Each DRIP enriched region was assigned to the genomic region for which it showed the greatest overlap. Enriched regions which displayed no overlap with any feature were assumed to locate within the Pol II PTUs. Enriched regions assigned to the Pol II PTUs were further classified as associated with the CDS or UTR sequences, or else assigned as intergenic. Motif analysis of Pol II PTU-associated enriched regions was done using MEME version 4.12.0 under default settings (77).

Normalised ratio bigwig DRIP-seq files were used to generate metaplots and heatmaps using deepTools (78). DRIP versus PAS metaplots were generating with the makemetaplot.pl script from HOMER (Heinz et al 2010) using PAS coordinates (61,79)and enriched region coordinates.

H3v, H4v, H2Av, H2Bz, H4k10ac and BDF3 ChIP-seq data was sourced from Siegel et al 2009, H3 ChIP-seq from Wedel et al 2017 and mRNA half-life from Fadda et al 2014. All data was processed as per publication methods.

### GC and AT skew analysis

GC and AT skew were calculated as (G-C)/(G+C) and (A-T)/(A+T), respectively. Skew was calculated either in 11 bp windows (analysis of ATG translational start sites only) or in 100 bp windows across the *T. brucei* hybrid genome. The results of whole genome analysis were converted into bigwig format and skew was plotted over regions of interest with deepTools analysis software.

## Results

### R-loops are highly abundant in the *T. brucei* genome

RNA-DNA hybrid immunoprecipitation (DRIP), either on cross-linked or ‘naked’ DNA, has been used in a number of organisms, with some experimental variations based on treatment of the nucleic acid components, the mode of genome fragmentation and whether the recovered nucleic acid was analysed by sequencing, microarray hybridisation or qPCR (80). Here, we used S9.6 for DRIP after formaldehyde fixation, an approach that has been applied to R-loop mapping in *Saccharomyces cerevisiae* (12,28,81,82), *Schizosaccharomyces pombe* (83) and *Arabidopsis thaliana* (84). To provide a genome-wide picture of R-loop distribution, DNA was isolated from the DRIP material and characterised by Illumina sequencing (DRIP-seq), mapping the reads to the assembled *T. brucei* genome (85). To ensure R-loops were mapped, DRIP-seq was performed in both wild type (WT) BSF *T. brucei* cells and in mutants lacking RNase H1 (TbRH1, encoded by gene Tb427.07.4930), allowing comparison of the mapping profiles. TbRH1 mutants were generated by two rounds of allele replacement, with the resulting *Tbrh1*−/− null cells found to be viable and to grow in culture at the same rate as WT BSF cells (Briggs et al, manuscript in preparation).

DRIP-seq mapping to the *T. brucei* genome revealed very widespread coverage (Fig.1): based on >1.2 fold enrichment of reads in the DRIP sample relative to input, ~35,000 enriched regions were predicted in both WT and *Tbrh1*−/− cells. To understand if the enrichment was localised to specific sequences, we divided the genome into regions transcribed by the three RNA Pols, largely non-transcribed VSG arrays and SSRs, and centromeric and retrotransposon and associated sequences (Fig.1A). DRIP-seq enriched regions were found in all these locations, but with notably highest abundance in the RNA Pol II polycistronic transcription units (PTUs; Fig.1A). Though loss of TbRH1 did not increase the number of enriched regions, greater abundance of putative R-loops were found in non-PTU regions (Fig.1A) and a greater amount of the genome was enriched (Fig.S1), consistent with spreading of R-loops following loss of the RNase H1 enzyme.

**Figure 1.**
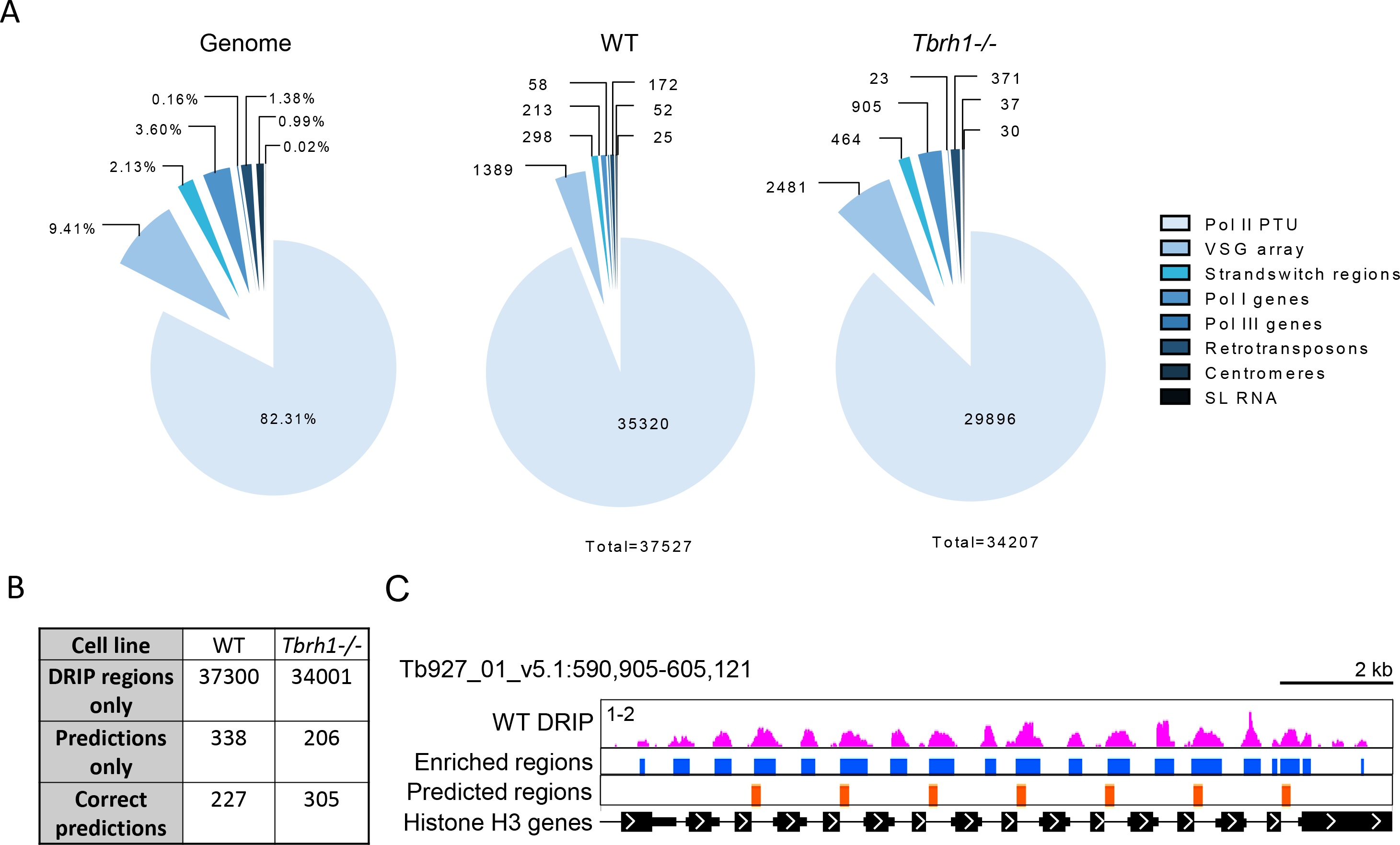
Distribution of R-loops in the genome of bloodstream form *T. brucei*. A) Analysis of R-loop locations in WT and *Tbrh1−/−* cells showing DRIP enriched region data sets (middle and right charts) relative to the sequence composition of the 11 *T. brucei* 11 megabase chromsomes (left chart); genomic elements are colour-coded according to right-hand key. B) Comparison of the number of WT and *Tbrh1−/−* DRIP enriched regions and predicted R-loop forming sequences. C) Screenshot of a histone H3-encoding region of chromosome 1. WT DRIP data is shown in pink (1-3 on y-axis denotes level of enrichment in DRIP relative to input), with identified enriched regions from this data set in blue, and predicted R-loop forming regions in orange. CDS are shown as thick black lines, UTRs as thin black lines and arrows show direction of transcription; size is indicated.

### R-loops are enriched at repeated sequences, including centromeres, RNA Pol I arrays and retrotransposon-associated genes

Given the large number of predicted DRIP-seq enriched regions, which massively exceed the number of *T. brucei* loci predicted to form R-loops (Fig.1B)(86), we examined the mapping in more detail to ask about read distribution. Within the RNA Pol II PTUs, DRIP-seq enrichment was clearly non-random, with a pronounced focus on regions separating predicted CDS (Fig.1C); this distribution is examined in more detail below. Comparing levels of DRIP-seq enrichment relative to predicted repeated sequences(87) further suggested a non-random distribution (Fig.S2), since many regions with higher levels of enrichment coincided with clusters of repeats. Amongst such repetitive regions were the *T. brucei* centromeres, which have been mapped to date in eight of the 11 diploid megabase chromosomes (88). DRIP-seq signal was strongly enriched at the A-T rich repeats found at the centromeres (Fig.2A) and levels of enrichment in all cases increased in the *Tbrh1*−/− mutants relative to wildtype (Figs2A,B; Fig.S3), indicating centromeres are a focus of R-loop formation. In common with other eukaryotes, *T. brucei* rRNA genes are transcribed by RNA Pol I and organised as gene arrays. Here, DRIP-seq enrichment was localised to the non-CDS regions in WT cells and the signal increased and spread in *Tbrhh1*−/− mutants (Fig.3A), indicating rRNA arrays are a conserved location of R-loops (24,81). In contrast with other eukaryotes, *T. brucei* protein-coding genes can also be transcribed by RNA Pol I, including procyclin and VSG. DRIP-seq enrichment was notably greater in the VSG expression sites (ES) that are actively transcribed in BSF *T. brucei* compared with the procyclin loci or metacyclic VSG expression sites (Fig.3A), from which gene expression is repressed in this life cycle stage (89). In addition, loss of TbRH1 caused a more pronounced increase in signal at the bloodstream VSG ES (Fig.3A), perhaps suggesting R-loops predominantly form co-transcriptionally at RNA Pol I loci. In yeast, DRIP-seq indicated R-loops form at RNA Pol III transcribed genes, and their abundance increases in yeast mutants lacking both RNase H enzymes (24). Here, DRIP-seq signal was enriched at both tRNA and snRNA genes but, paradoxically, enrichment was decreased in the *Tbrh1*−/− mutants (Fig.3B). A similar DRIP-seq profile was revealed for snoRNA gene arrays (Fig.S4), which are probably transcribed by RNA Pol II (90), suggesting not all repeated or structural *T. brucei* RNAs form R-loops that are acted upon by RNase H1. Finally, DRIP-seq enrichment revealed dramatically increased signal at retrotransposon hotspot (RHS) genes in *Tbrh1*−/− cells relative to WT (Fig.S5). RHS genes comprise a highly abundant, variable gene family in *T. brucei* and *T. cruzi* (91,92). Members of the RHS family express nuclear proteins and are frequent targets for transposable elements, though their functions remain elusive. Thus, though DRIP-seq enrichment here may be related to transposable elements being targets for R-loop formation in yeast (24) and plants (51), RHS localisation may indicate kinetoplastid-specific R-loop functions.

**Figure 2.**
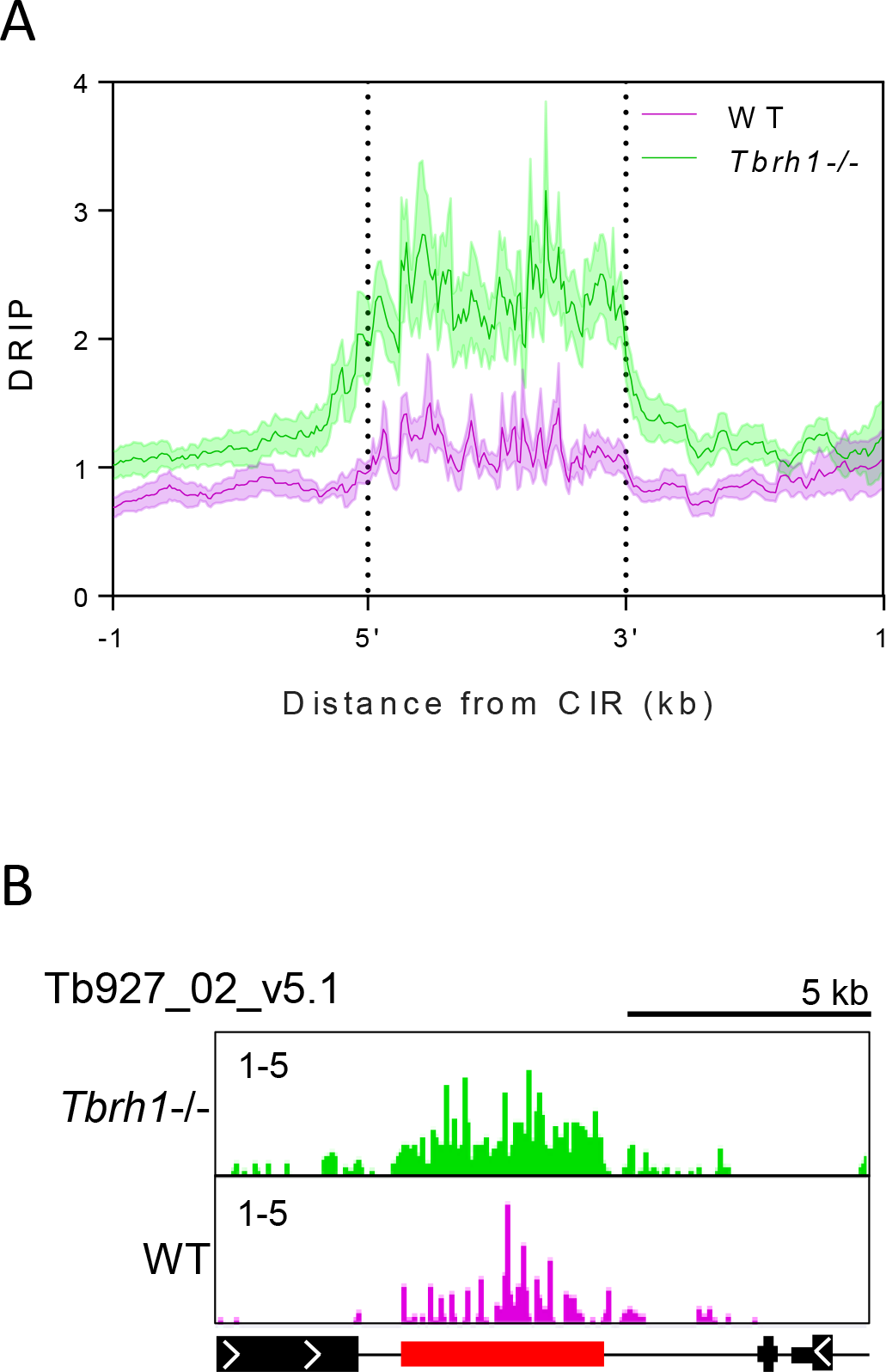
R-loops are enriched at *T. brucei* centromeres. A) Metaplot of DRIP signal in WT (pink) and *Tbrh1−/−* cell (green) data sets centred on the annotated centromeric interspersed repeats (CIR) +/− 1 kb. B) Representative screenshot of the a portion of chromosome 2 (Tb927_02_v5.1) containing the centromere region; CDS and DRIP-seq enrichment annotations are as shown as in Fig. 1, CIR are shown in red.

**Figure 3.**
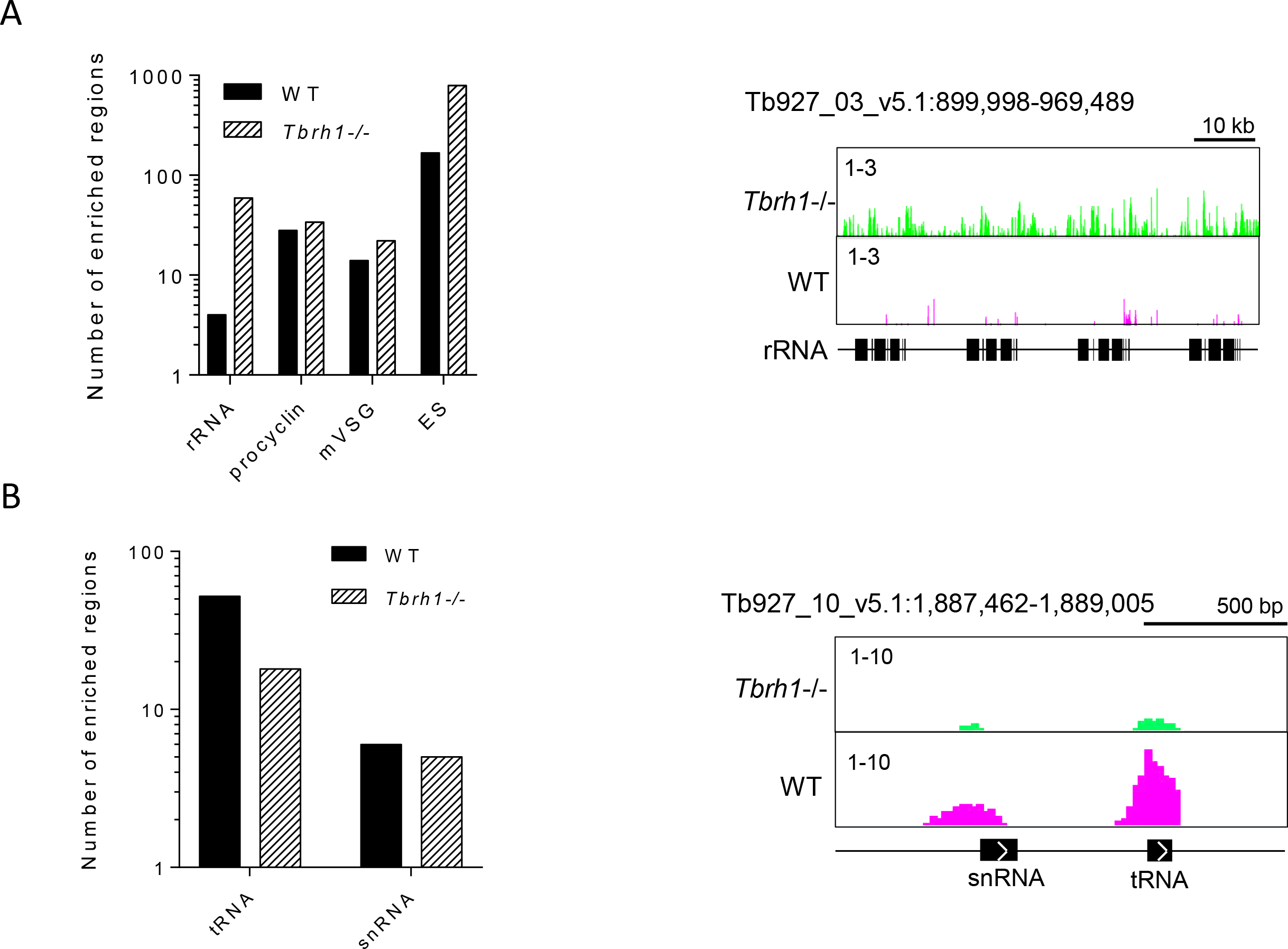
DRIP-seq signal at RNA Pol I and Pol III transcribed sites in *T. brucei*. A) Graph showing the number of enriched regions identified at Pol I transcribed loci for WT and *Tbrh1−/−* data sets (left); a representative screenshot of a rRNA locus is shown (right; annotation as in Fig.1). B) Graph showing the number of enriched regions identified at Pol III transcribed loci (left) and a representative screenshot of a locus containing snRNA and tRNA genes (right, annotation as in Fig.1).

### Most *T. brucei* R-loops co-localise with sites of multigenic transcript processing

As discussed above, DRIP-seq enrichment was most abundant throughout the RNA Pol II PTUs, with signal in WT cells mainly confined to intra-CDS sequences. Such localisation was not limited to repeated genes, such as histone H3 (Fig.1C), but was true for all RNA Pol II transcribed genes. In WT cells and in *Tbrh1*−/− mutants 34% and 39%, respectively, of enriched regions localised to CDS (Fig.4A), despite these sequences comprising 53% of the PTUs. Plotting of DRIP signal over genes that appeared to have intra-CDS DRIP-seq enrichment and those that do not showed little difference in DRIP profile, indicating intra-CDS R-loops are weakly enriched compared to those at the flanking sites (Fig.S7A). Further examination of the intra-CDS R-loop positive genes did not reveal a clear pattern: base composition, CDS length, mRNA half-life and UTR length appeared indistinguishable from genes without intra-CDS R-loops (Figs.S7B,C); and, for protein-coding genes, no clearly enriched GO terms could be found in the cohort (Table S1). Thus, unlike in *A. thaliana* (51) R-loops in *T. brucei* appear to be mainly excluded from RNA Pol II-transcribed coding sequence and, when present there, cannot easily be assigned a function.

**Figure 4.**
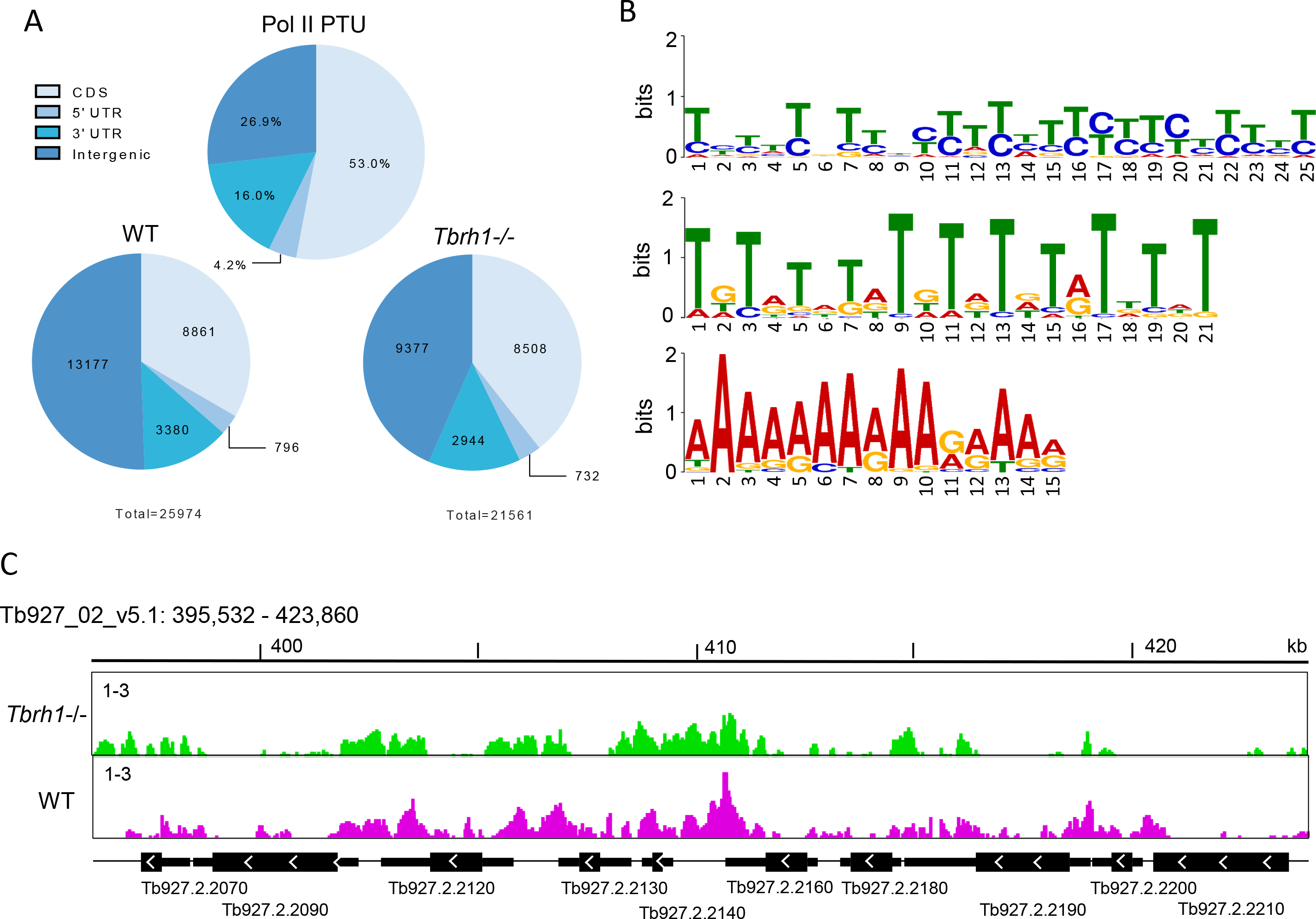
R-loops within RNA Pol II polycistronic transcription units are predominantly intergenic. A) Analysis of R-loop locations in WT and *Tbrh1−/−* cells showing DRIP enriched region data sets within the Pol II PTUs (lower charts) relative to the sequence composition of Pol II PTUs (upper chart); regions are annotated according to the key. B) Three motifs identified by MEME analysis of WT enriched regions localised to within the Pol II PTUs. C) Representative screen shot of WT and *Tbrh1−/−* DRIP data over a region (Tb927_02_v5.1: 395,532 - 423,860) within a Pol II PTU in chromosome 2. Gene and DRIP-seq enrichment annotations are as shown in Fig.1.

Motif analysis using MEME revealed enrichment within Pol II PTU-associated R-loops for three interesting motifs; two polypyrimidine sequences and one poly(A) tract (Fig.4B), suggesting R-loops localise to sequences associated with the *trans* splicing and polyadenylation events needed to generate mature mRNAs from multigene RNAs. Indeed, detailed mapping confirmed the predominance of DRIP-seq reads in the intergenic sequences and untranslated regions (UTRs) in both WT and *Tbrh1*−/− cells (Fig.4C, Fig.S6). Heatmaps of DRIP-seq enrichment around every RNA Pol II gene revealed relatively precise signal localisation upstream and downstream of the CDS for probably all genes and showed that enrichment increased after loss of TbRH1 (Fig.5A). To understand this localisation we first asked if the DRIP-seq enrichment pattern correlated with mapped sites of trans-splicing and polyadenylation (61,93), as suggested by MEME analysis. The density of polyadenylation sites (PAS) and DRIP enriched regions showed a remarkably strong correlation when analysed as density relative to distance upstream and downstream of CDS in both WT and *Tbrh1*−/− mutants (Fig.5B). Visualisation of the mapped reads confirmed this association, with levels of DRIP-seq enrichment notably higher in regions of clustered PAS and with patterns that follow the direction of transcription and distance from the upstream CDS (Fig.5C, Fig.S8), suggesting R-loops form at these sites during RNA processing. In contrast, DRIP-seq enrichment showed a less clear association with mapped splice acceptor sites (SAS) (Fig.5C), though it might be noted that these are less abundant than PAS in inter-CDS sequence.

**Figure 5.**
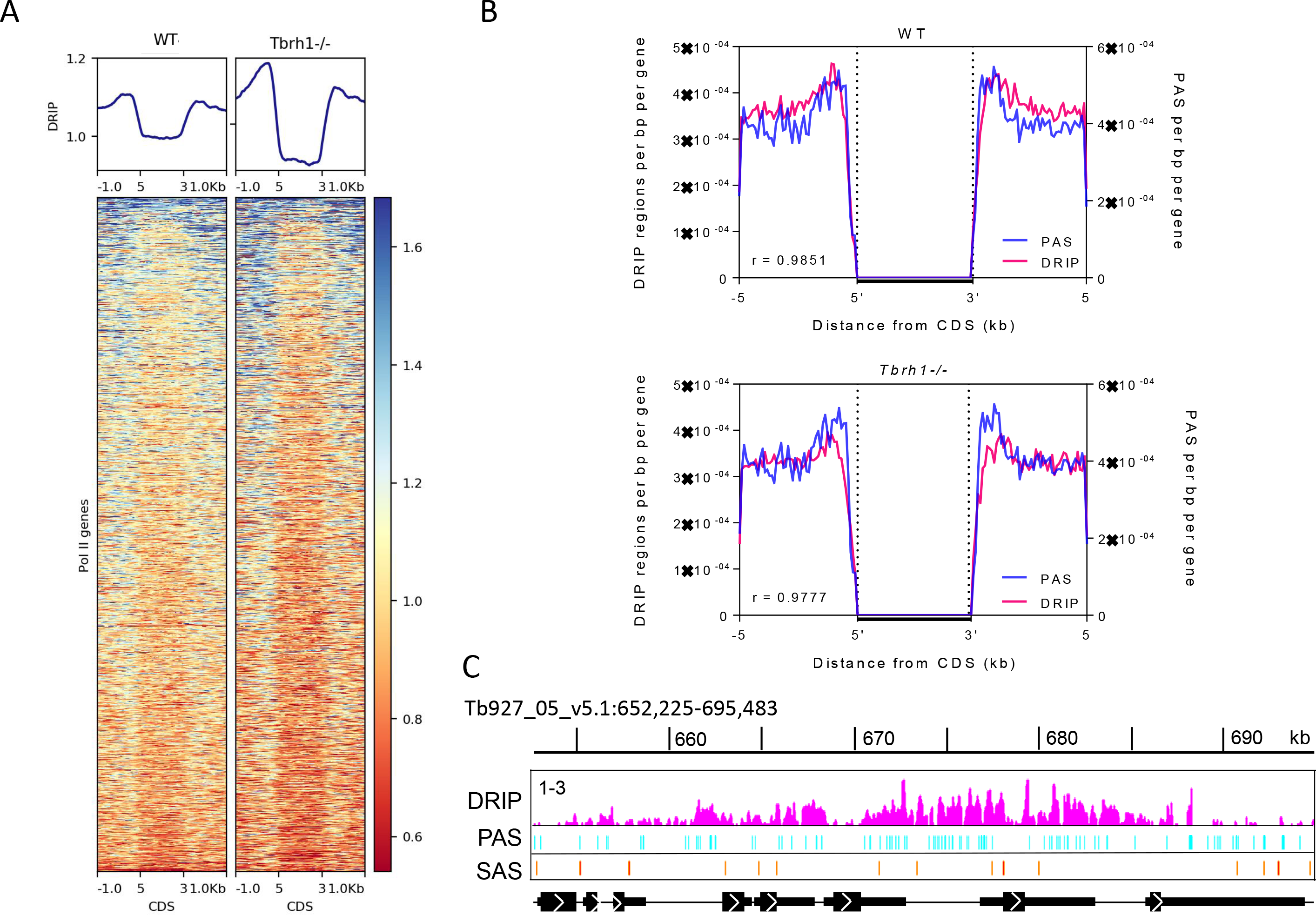
R-loop accumulaton in RNA Pol II transcription units strongly associate by polyadenylation sites. A) Metaplots and heatmaps of WT and *Tbrh1−/−* DRIP signal over the CDS +/− 1 kb of each Pol II transcribed gene. B) Metaplots of the number of UTR-or intergenic-associated DRIP enriched regions (red) and PAS (blue) regions per bp for each Pol II CDS +/− 5 kb for WT (upper) and *Tbrh1−/−* (lower) DRIP-seq data. C) Prepresentative screenshot of WT DRIP signal (pink) relative to mapped PAS (blue) and SAS (orange) locations in a region of chromosome 5; CDS and DRIP-seq enrichment annotations are as shown as in Fig. 1.

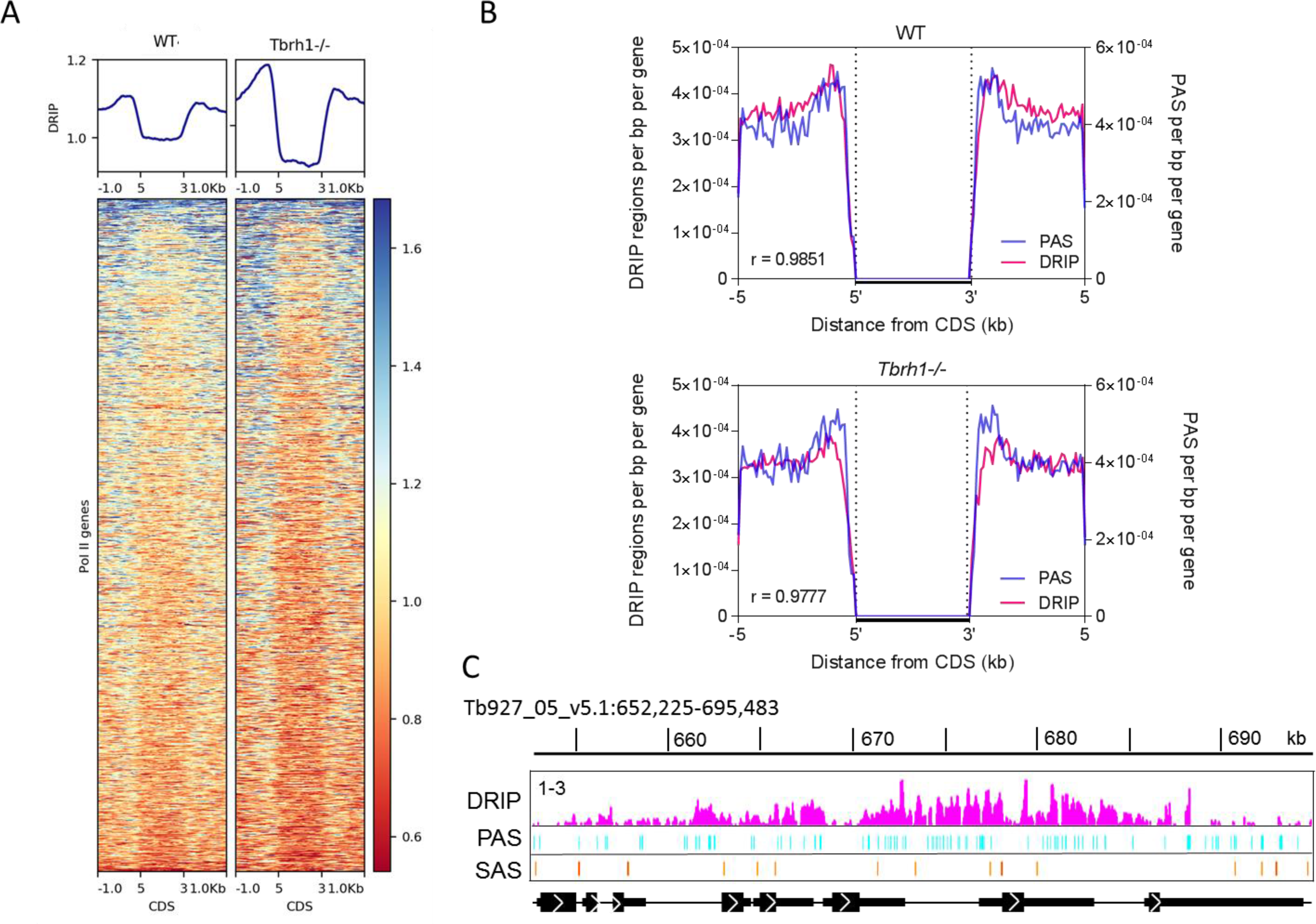

Next, to examine R-loop positioning relative to CDS, we plotted the level of DRIP-seq enrichment relative to the start codon of every RNA Pol II transcribed gene. To do this, we separated the genes into those predicted to be first within a PTU (n, 110), and therefore proximal to the transcription start sites, and all others (n, 9977), which are distal to the transcription start site and internal to the PTU. In both cases a pattern of DRIP-seq enrichment was revealed (Fig.6A) in which read abundance peaked upstream of the ATG and displayed a region of depleted reads downstream of the ATG. Though in WT cells this pattern was more pronounced for PTU-internal genes compared with the first genes in the PTU, in both cases the level of DRIP-seq signal increased and the pattern became more pronounced in *Tbrh1*−/− mutants (Fig.6A), most notably upstream of the first gene ATG sites were transcription begins. In both cases, therefore, the DRIP-seq detected R-loops. Very recently, histone H3 occupancy has been described around RNA Pol II transcribed genes in *T. brucei*, revealing nucleosome depletion upstream of every ATG (63). Mapping the H3 ChIP-seq of Wedel et al alongside our DRIP-seq data (Fig.6B) revealed a striking correspondence: for both the first genes of the PTUs and internal PTU genes the patterns of nucleosome depletion and R-loop accumulation upstream of the ATG closely mirrored each other; moreover, the region of R-loop depletion downstream of the ATG appeared to follow a small peak of increased nucleosome abundance. Taken together, these data reinforce the association of R-loops in *T. brucei* with sites of RNA processing and suggest RNA-DNA hybrids form at locations of ordered nucleosome positioning that might influence RNA Pol II movement to facilitate trans-splicing and polyadenylation. Examination of the AT and GC content of the sequences around the ATGs relative to R-loop enrichment (Fig.S9) suggested some increase in GC skew as sequences become more distal upstream and downstream of the ATG, a pattern that was more marked when analysing the greater number of PTU-internal genes. Interestingly, slight GC and AT skew was found to be inversely correlated to WT DRIP signal around the ATG of all Pol II genes (Fig.S9), which contrasts with findings that R-loops are associated with positive GC skew at human CpG island promoters (47) and both AT and GC positive skew is associated with R-loops in the *Arabidopsis* genome (51).

**Figure 6.**
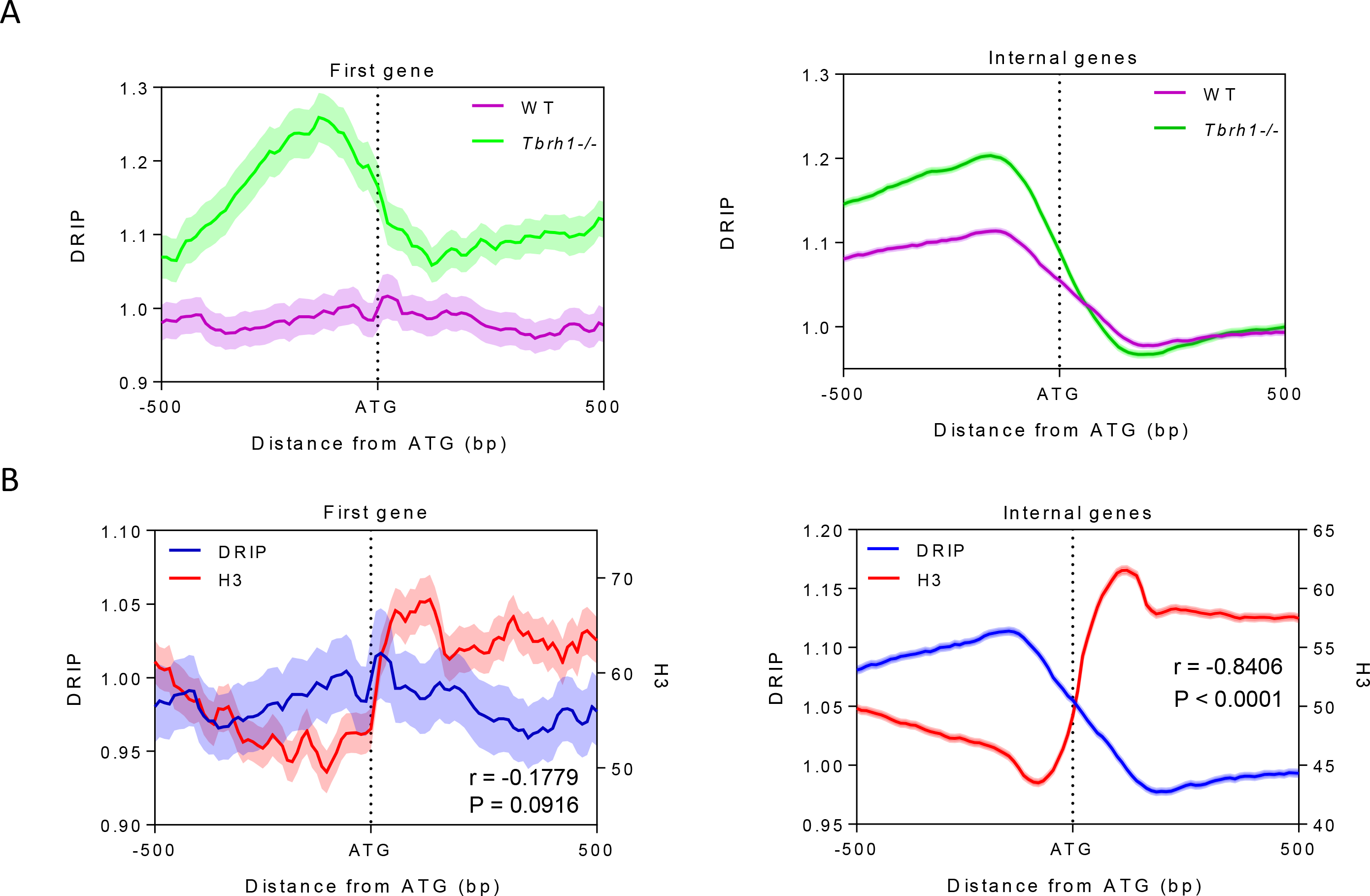
R-loop accumulation is the mirror of nucleosome accumulation and is dictated by RNaseH1 throughout RNA Pol II transcription units. A) Metaplots of WT (pink) and *Tbrh1−/−* (green) DRIP signal over the ATG (+/− 500 bp) of the first gene of each Pol II PTU (left) and over all other genes in the PTU (right). B) Metaplot analysis of DRIP (blue) and histone H3 ChIP (red) signal over the ATG (+/− 500 bp) of the first gene of each Pol II PTU (left) and all other Pol II transcibed genes (right).

### No evidence for R-loop localisation at *T. brucei* DNA replication origins

Sites of DNA replication initiation, termed origins, have been mapped in a subset of SSRs in *T. brucei* by MFA-seq (67,68). What features distinguish these origin-active SSRs from origin-inactive SSRs is unclear, since one component of the *T. brucei* Origin Recognition Complex, TbORC1/CDC6, appears to map to all SSRs (69,94). In addition, no sequence features have been described that distinguish the two classes of SSRs, with the exception of highly active origins being coincident with centromeres in at least eight chromosomes (70). To ask if R-loops might represent a hitherto undetected epigenetic feature that directs origin activity, we separated SSRs into those in which replication initiation has been mapped by MFA-seq and those in which origin activity has not been detected, and examined the patterns of DRIP-seq enrichment (Fig.7). Irrespective of whether the SSRs were origin-active (Fig.7A) or -inactive (Fig.7B) a similar pattern of DRIP-seq enrichment was seen, with a striking depletion in signal around the centre of the SSR and increased signal approaching the most proximal genes. MFA-seq is unable to determine if DNA replication initiates at discrete sites within an SSR or is dispersed throughout the loci. However, as the DRIP-seq signal showed comparable levels of enrichment at the centre and CDS-proximal sites of origin-active SSRs compared with -inactive SSRs, and the DRIP-seq enrichment increased in both types of SSRs in *Tbrh1−/−* mutants relative to WT cells, it seems clear that though R-loops form within SSRs they show no differential localisation that could explain the differing patterns of DNA replication initiation at these inter-PTU loci.

**Figure 7.**
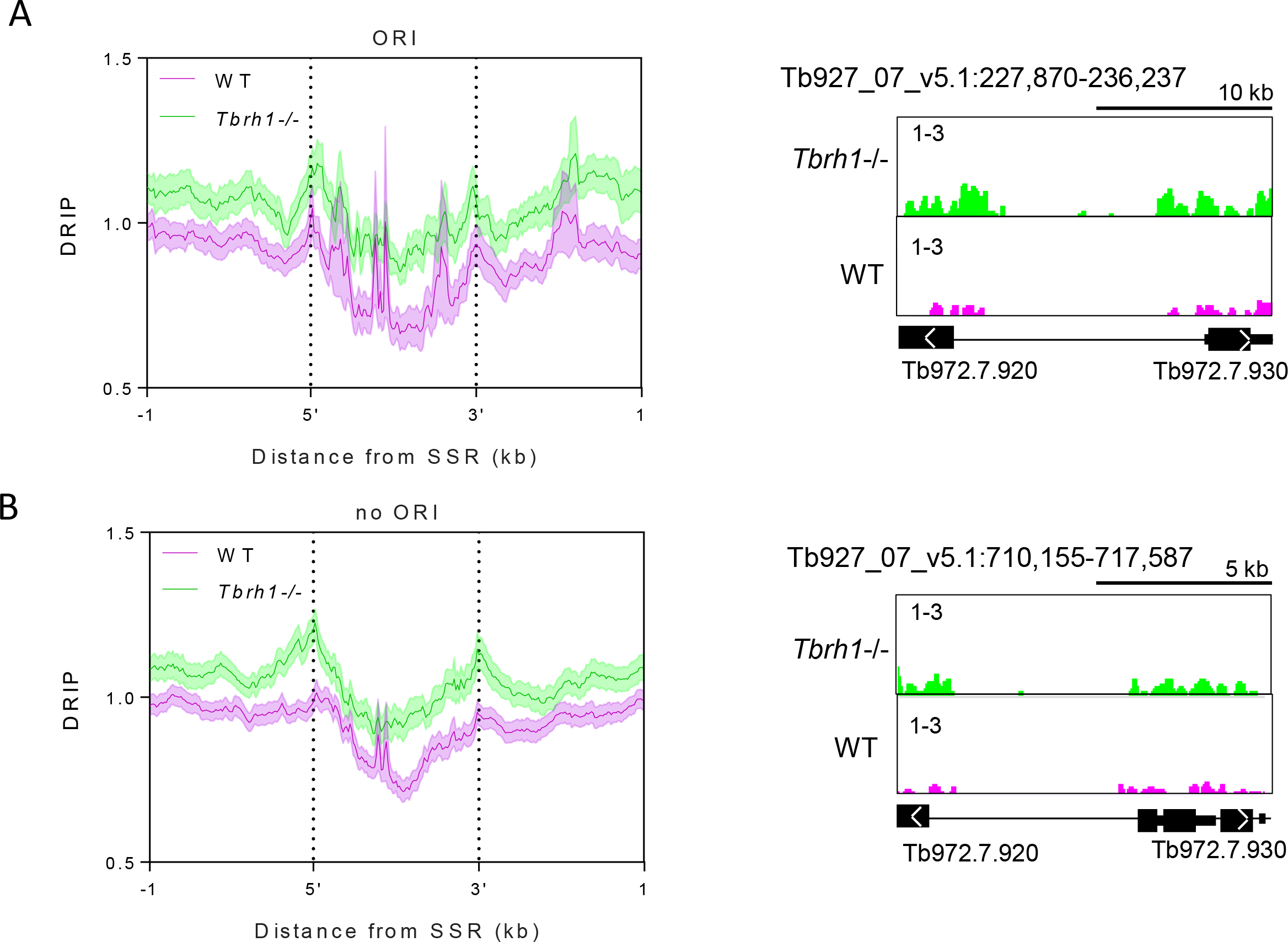
R-loop distribution is equivalent at strand switch regions at which replication initiation has been mapped, or where replication initiation has not been detected. A) Metaplot analysis of WT (pink) and *Tbrh1−/−* (green) DRIP signal over replication origin (ORI)-associated SSRs (left), with a preresentative screenshot of one such SSR (right; gene and DRIP-seq enrichment annotations are as shown in Fig.1). B) Metaplot (left) and representative screenshot (right) of SSRs where no replication origin (no ORI) activity has been detected.

### R-loops are enriched at sites of transcription initiation

The above data suggest increased enrichment of R-loops in regions of the SSRs that are transcribed, or in elements that direct transcription initiation or termination. In other eukaryotes, R-loops have been localised to regulated promoters, suggesting roles in controlling transcription initiation (40–43), and to the ends of genes, suggesting roles in transcription termination (44–47). To ask if such R-loop functions might also act in *T. brucei*, we separated the SSRs into divergent, convergent or head-to-tail loci, where transcription and histone mapping predicts, respectively, initiation of transcription at divergent PTUs, termination of transcription at convergent PTUs, and mixed sites with transcription initiation of one PTU and termination of another. Evaluating DRIP-seq enrichment patterns in the three classes of *T. brucei* SSR suggests a role in transcription initiation but with less clear evidence for a role in termination (Fig.8). In divergent SSRs (Fig.8A) two pronounced peaks of DRIP-seq enrichment were found at the boundaries of the loci, both upstream of the first predicted genes of the two divergent PTUs. In head-tail PTUs (Fig.8B), a strong peak was again seen upstream of the first gene in the PTU, but a peak of similar magnitude was not seen around the final gene of the other PTU. In both types of SSR the level of DRIP-seq signal increased in the peaks adjacent to the transcription start sites in *Tbrh1−/−* cells relative to WT, indicating the peaks represent R-loops. Though DRIP-seq peaks could be discerned in convergent SSRs (Fig.8C), this pattern was the result of high signal enrichment at tRNA genes in 2 of the 15 convergent SSRs analysed. Since the same pattern was also seen downstream of the PTUs in some head-tail SSRs (Fig.8B), and the level of DRIP-seq enrichment at the ends of the PTUs in both types of SSR was not increased in *Tbrh1*−/− mutants compared with WT cells (Fig.8B and C), DRIP-seq provides weaker evidence for R-loop association with sites of multigenic transcription termination compared with sites of initiation.

**Figure 8.**
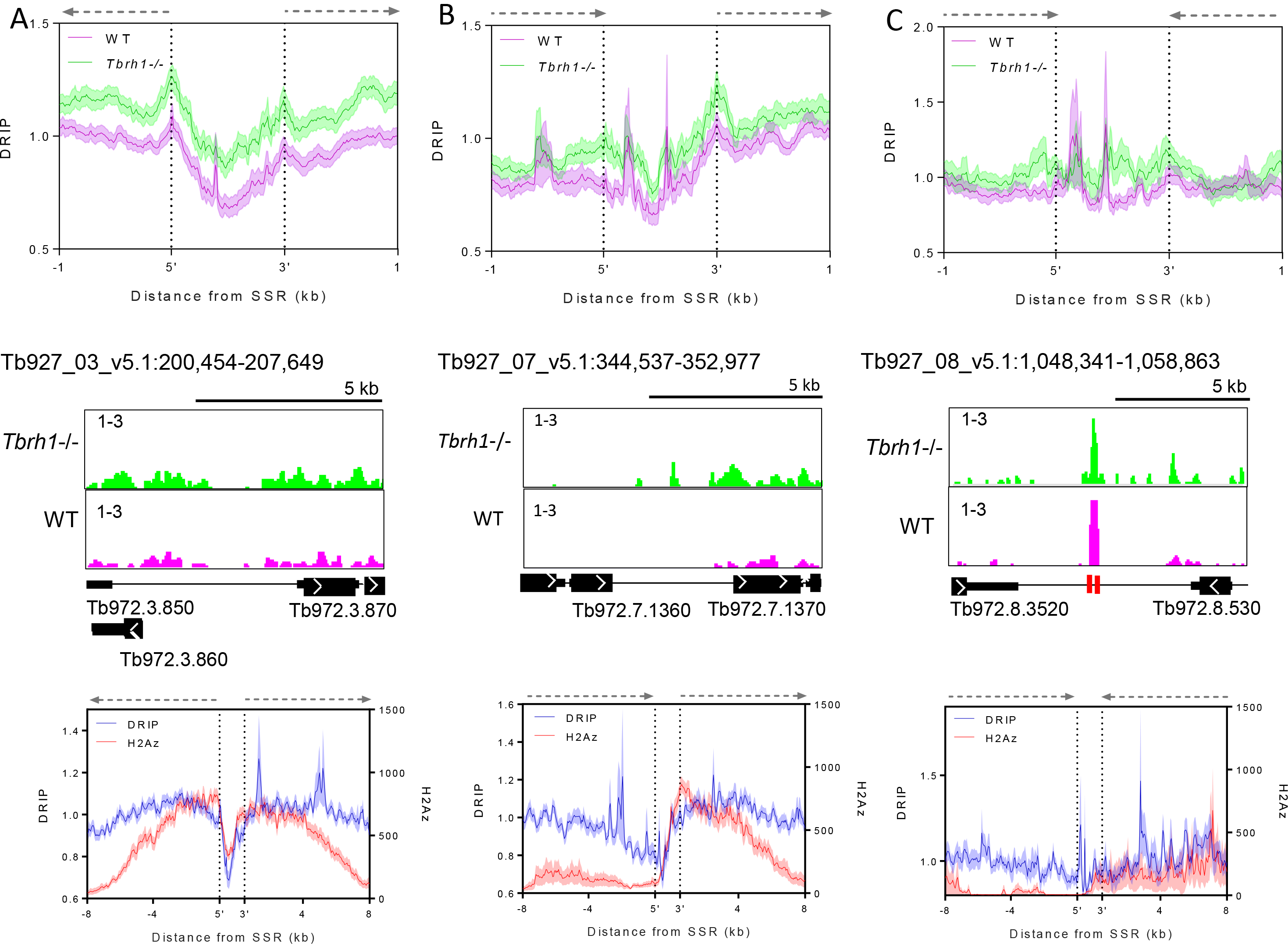
R-loop enrichment shows a greater association with multigenic transcripton initiation and associated markers than termination. The upper diagrams show metaplot profiles of WT (pink) and *Tbrh1−/−* (green) DRIP signal over divergent (A), head-to-tail (B), and convergent (C) SSRs (+/− 1 kb), and the middle diagrams are representative screenshots of each (gene and DRIP-seq enrichment annotations are as shown in Fig.1). In the lower diagrams the WT DRIP signal (blue) metaplot is compared to H2Az ChIP signal (62) over divergent (left), head-to-tail (middle) and convergent (right) SSRs (+/− 8 kb).

To further examine the association of R-loops with transcription initiation we generated metaplots of the WT DRIP-seq data at each class of SSR, extending the analysis to 8 kb upstream and downstream of the SSR boundaries, and comparing the profiles to that of previously published ChIP-seq data sets (62) of the following factors associated with transcription initiation: histone variants H2Bv and H2Az, histone H4 acetylated on lysine 10 (H4K10ac) and bromodomain factor BDF3 (Fig.8, FigS10). These plots revealed strong correlation of H2Az, H2Bv and H4K10ac signal with DRIP signal at the boundaries of divergent SSRs, though the correlation diminished when the plots extended into the PTUs, since here R-loops are present but enrichment of the variant and modified histones is not seen (62). The same correspondence was also seen at the transcription initiation region of head-to-tail SSRs (unlike at the termination region; see below). Enrichment of BDF3 did not correlate as well with the DRIP-seq signal, as enrichment of the bromodomain factor peaked within the SSR boundaries, and therefore upstream of the highest point of DRIP enrichment, at divergent SSRs (a pattern also seen at the 3’ boundary of head-to-tail SSRs; Fig.S10). Within convergent SSRs, as well as at the ends of PTUs in head-to-tail SSRs, there was a lack of enrichment of all four factors, reflecting the limited enrichment of DRIP-seq signal. In contrast, enrichment of the transcription termination-associated histone variants H3v and H4v was notably greater than that of DRIP-seq(Fig.S11), further indicating R-loops display less association with transcription termination compared with initiation in *T. brucei*.

## Discussion

Work in a range of eukaryotes, including plants, insects, yeast and mammals, is revealing widespread functional roles for RNA-DNA hybrids termed R-loops. Prominent amongst these emerging activities are roles for R-loops in the control of RNA Pol II transcription initiation and termination, as well as wider localisation to repeated sequences and non-coding RNA. R-loops can also cause genome instability, chromatin modifications and have been implicated in DNA replication. Much of the observations on R-loop activities are based on characterisation of ‘conventional’ eukaryotic protein-coding genes, each of which is a self-contained unit bounded by a dedicated upstream RNA Pol II promoter and, downstream, by a termination region. Indeed, in some cases R-loops have been functionally associated with well understood promoters whose activity is regulated by specific RNA Pol II transcription factors. Here we have mapped R-loop localisation in the highly unconventional genome of *T. brucei*, revealing potentially diverged roles for the RNA-DNA hybrids in the mRNA processing steps needed during multigene transcription. In addition, we find pronounced R-loop localisation at poorly defined and putatively unregulated RNA Pol II transcription initiation sites. Finally, we reveal conserved roles for *T. brucei* R-loops, including localisation to transcribed and untranscribed repeated genes and sequences.

Virtually every RNA Pol II transcribed gene in *T. brucei* is encoded from a multigenic transcription unit, which has resulted in the evolution of widespread and coupled trans-splicing and polyadenylation to separate adjacent genes into mature (capped and polyadenylated) mRNAs (56). The near genome-wide use of RNA Pol II multigenic gene expression is common to all kinetoplastids and its extent has no known parallel in other eukaryotic groupings. DRIP-seq mapping revealed that the most prevalent localisation of R-loops in the *T. brucei* genome is within the intergenic regions between CDS in the multigenic transcription units. Though R-loops could be detected within some CDS, this was a minor region of DRIP-seq enrichment, which contrasts with the more equitable distribution of R-loops between gene bodies and intergenic regions observed in *A. thaliana* (51). Intra-CDS R-loops have also been described in *S. cerevisiae*, strongly correlating with highly expressed genes (52). In *T. brucei*, we have been unable to identify gene features, including mRNA half-life, mRNA length or predicted function, which dictate intra-CDS R-loop accumulation. Importantly, it is very unlikely that transcription rate can explain intra-CDS R-loops, since multigenic transcription would suggest each gene within a PTU is covered by equivalent densities of RNA Pol II. Whether sequence features can alter the dynamics of RNA Pol II movement across select genes, or if some mRNAs are present for longer in the nucleus before export, allowing for increased DNA interaction, is unknown (64). Given these limitations, it is unclear if intra-CDS R-loops might function to influence gene expression. In contrast, the pronounced localisation of R-loops between CDS and throughout the Pol II PTUs suggests the major genomic enrichment of R-loops in *T. brucei* is a novel association with RNA processing, reflecting the multigenic nature of kinetoplastid transcription (Fig.9).

**Figure 9.**
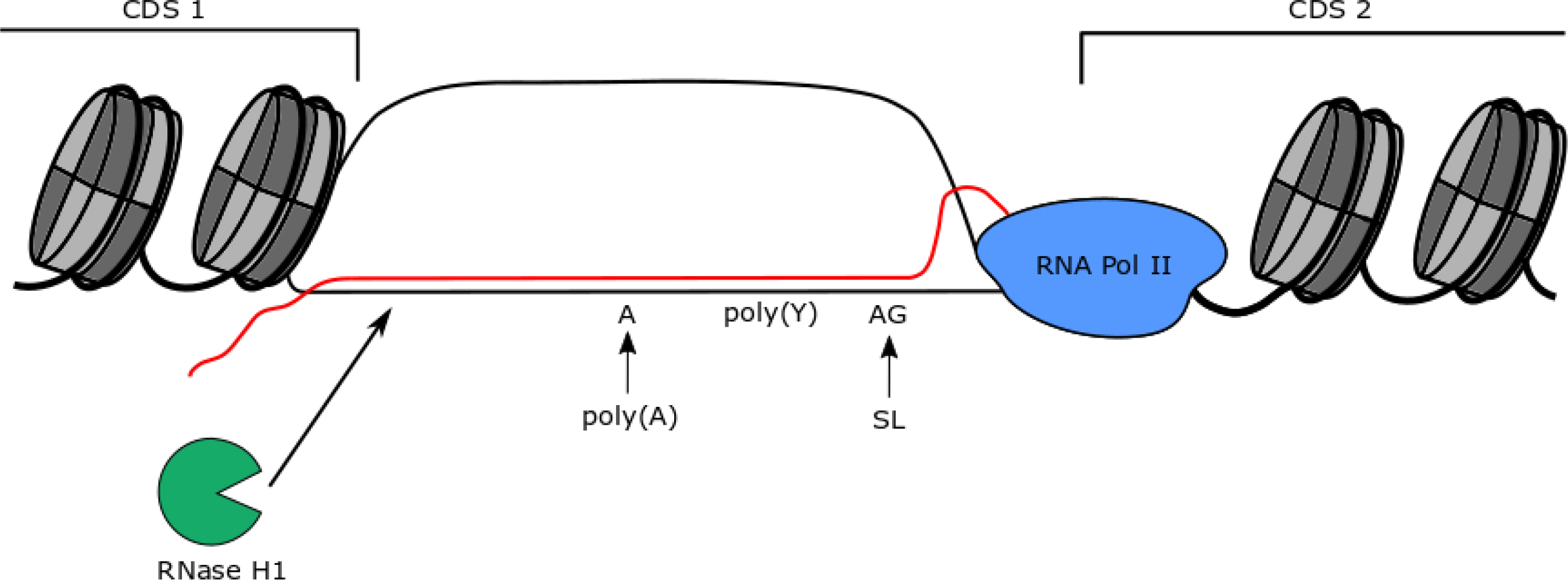
A summary of R-loop localisation at mRNA processing regions in *T. brucei* multigenic RNA Pol II transcription units. Genes within Pol II PTUs are separated by UTR and intergenic DNA sequences containing sites of polyadenylation (poly(A)) and splice acceptor addtion (AG, where the splice leader (SL) 5’ cap is trans spliced), which are known to be directed by polypyrimidine (poly(Y)) tracts that are critical for correct maturation of the mRNA. R-loops containing RNA (red) putatively emerging from RNA Pol II (blue) and hybridising to the DNA (black)form over these regions between the CDS of adjacent genes, within areas of nucleosome (grey) depletion, and are acted upon by *T. brucei* RNaseH1 (green).

In contrast to other eukaryotes, 5’ caps are added to kinetoplastid mRNAs by addition of a common spliced leader RNA through trans-splicing of multigene precursor RNA (95). Mechanistically trans-splicing is related to the removal of introns in other eukaryotes by cis-splicing, with each gene being in effect an exon. Less is understood about polyadenylation of kinetoplastid mRNAs (96,97), but the reaction follows trans-splicing and localisation of the polyA tail is guided by a polypyrimidine tract and AG sequence upstream of ORFs that predominantly dictate splicing (98,99). DRIP-seq reveals abundant R-loops within the intergenic regions associated with trans-splicing and polyadenylation, with a strong correlation between R-loop enrichment and polyadenylation site density and proximity to the gene CDS. MEME analysis also revealed enrichment of the expected polypyrimidine tractswithin Pol II PTU associated R loops, though why a poly(A) tracts was also identified is less clear, since such a feature has not been linked with trans-splicing or polyadenylation in PTUs. These data reveal a previously undescribed association between R-loops and mRNA processing (Fig.9). In yeast and humans, mapping of R-loops shows their abundance and density to be lower in intron-rich genes compared with genes lacking introns, at least in part because recruitment of the splicing machinery suppresses R-loop formation (100,101). In addition, mapping catalytically inactive RNase H1 in human cells indicates that in the rare instances where R-loops are detected within genes or at poly(A) sites, they may result from RNA processing errors and do not obviously correlate with RNA Pol II pausing sites (102). Taken together, these observations underline the novelty we see in *T. brucei*, where the predominant localisation of R-loops is within RNA Pol II-transcribed multigenic units at sites where RNA processing is known to occur. Importantly, localisation of R-loops to sites of polyadenylation in *T. brucei* is very unlikely to be related to the association seen in other eukaryotes between R-loops and RNA Pol II transcription termination (44–47), since at all such sites within the *T. brucei* PTUs termination must be avoided to ensure expression of downstream genes. Whether these data mean R-loops in *T. brucei*, unlike in eukaryotes with more conventional RNA Pol II genes, actively participate in the generation of mature mRNAs from a multigene precursor RNA is unclear, especially given the paucity of our understanding of polyadenylation. For instance, is it possible that RNase H cleavage of the RNA within the R-loop allows access to the polyadenylation machinery (97)? The non-essentiality of TbRH1 argues against such a crucial role, but this activity might be mainly assumed by RNase H2, which has not to date been functionally examined in *T. brucei*, or even shared between the two enzymes. An alternative explanation for R-loop localisation within *T. brucei* PTUs might lie in the precise correlation between regions of R-loop accumulation and nucleosome depletion upstream of the ATG of all genes. This correlation may indicate RNA-DNA hybrids form at sites where RNA Pol II pauses around a well ordered nucleosome (63), in which circumstances the RNA extruded from the Pol may have greater time to bind DNA. While this does not discount the possibility that R-loops actively participate in mRNA generation, it is possible the pattern of DRIP-seq enrichment within the PTUs reflects the dynamics of RNA Pol II travel due to the co-ordination of mRNA processing. Indeed, ordered nucleosomes at the 5’ end of exons have been proposed to play the same role in slowing RNA Pol II and promoting cis-splicing in other eukaryotes (63,103). However, such a functional correspondence between chromatin structure and trans-splicing or cis-splicing cannot be readily reconciled with the strong positive correlation between *T. brucei* R-loops and mRNA processing sites compared with the poor correspondence between R-loops and cis-splicing or RNA Pol II pausing in yeast and humans (100–102). Thus, R-loop mapping revealed by DRIP-seq in *T. brucei* may reflect genuine novelty in RNA biology as a result of multigenic transcription.

SSRs separating adjacent PTUs are the sites of multigenic RNA Pol II transcription initiation and termination in *T. brucei* and all kinetoplastids. Given the precedence of R-loops contributing to both processes in other eukaryotes (8,10), DRIP-seq mapping in these loci was revealing, despite the relative lack of functional characterisation of kinetoplastid RNA Pol II promoters and terminators. DRIP enrichment was most pronounced within *T. brucei* SSRs around the first gene of a PTU, indicating an association with transcription initiation. Promoters of metazoans have been separated into different classes based on transcription start site profile (104): regulated promoters have well defined start sites and mainly lack CpG islands, whereas dispersed start sites are found at promoters with CpG islands. Dispersed promoters can be ubiquitously expressed in all cell types or display developmental regulation. Though dispersed promoters are also found in *S. pombe* (105) they are less common than in metazoans, perhaps due to lack of CpG islands. In *T. brucei*, discrete transcription start sites within SSRs appear absent (61), suggesting similarities with metazoan dispersed promoters (63), though no CpG islands or examples of transcriptional control have been detailed. R-loops have been described downstream of transcription start sites and are prevalent in human genes containing CpG islands (40,41,47), with the RNA-DNA hybrids in some at least cases contributing to the control of gene expression. R-loops have also been implicated in controlling expression from the *A. thaliana* FLC locus (43), and common features of R-loop association with metazoan transcription initiation are histone modifications and non-coding RNA. In contrast, R-loops are less clearly associated with promoters in yeast (24,28). DRIP-seq enrichment in *T. brucei* SSRs, in association with variant and modified histones, around the first PTU gene suggests R-loops might be broadly connected to transcription initiation at dispersed promoters and do not merely provide a means of transcriptional activation control. Nonetheless, the basis for R-loop deposition in *T. brucei*, and whether it is conserved with metazoans, is unclear: to date non-coding RNAs within *T. brucei* SSRs have only been described at sites where two PTUs converge (terminate)(106); the epigenetic marks so far localised to SSR transcription initiation sites (variant histones H2AZ and H2BV, and histone H4K10Ac)(62) do not overlap with the chromatin signatures described at promoter-associated R-loops in humans and *A. thaliana*; and our data suggest it is not obvious that positive GC skew dictates *T. brucei* R-loop formation. In addition, the localisation of R-loops within *T. brucei* SSRs, and therefore upstream of, or close to, transcription initiation, may be distinct from the predominant localisation of metazoan R-loops downstream of transcription initiation. Thus, whether or not R-loops in *T. brucei* can play a role in directing transcription initiation by RNA Pol II is unclear; if R-loops play such a role, the non-essentiality of TbRH1 argues that this RNase H does not provide a critical function.

In contrast to the pronounced association of *T. brucei* R-loops with initiation of multigenic transcription, the DRIP-seq data provides less compelling evidence for a role in termination, since signal enrichment at the ends of PTUs, such as within convergent SSRs, is less pronounced. In addition, where peaks are seen at such sites they frequently localise to tRNAs and do not become more pronounced in *Tbrh1*−/− mutants (unlike R-loop abundance at the start of PTUs). The limited enrichment we describe contrasts with the strong localisation of R-loops at gene termination sites in other eukaryotes (44–47). This difference between *T. brucei* and characterised eukaryotes is perhaps surprising, since non-coding RNAs have been implicated in R-loop action during termination (44,107), and such RNAs are readily detected where multigenic transcription units converge in *T. brucei* (106). Conceivably, factors other than TbRH1, such as helicase (45,108) or cleavage complex (46) orthologues, could provide a greater role in resolving R-loops at termination sites within SSRs, meaning the DRIP-seq approach adopted here may have missed a conserved termination mechanism. However, a novel mechanism remains possible given the emerging role of base J (a modified base found only in kinetoplastids and close relatives) in RNA Pol II transcription termination (65,66,109,110).

Localisation of R-loops to potentially all *T. brucei* SSRs at which transcription initiation occurs appears to rule out a role for the RNA-DNA hybrids in directing initiation of ORC-directed DNA replication, since the available mapping data suggest that only a fraction of SSRs are used constitutively as origins (67,68). In addition, origins have been described at convergent SSRs, where we see less evidence for R-loops. It is possible that R-loops contribute to recruitment of the *T. brucei* origin recognition complex, as proposed in humans (39), since ORC1/CDC6 is not limited to origin-active SSRs (67); indeed, loss of ORC1/CDC6 alters RNA levels at the ends of the PTUs, perhaps reflecting links between R-loops, replication and transcription. Nonetheless, discrete R-loop association with specific SSRs appears unable to explain why DNA replication strongly initiates at only some of the SSRs, an observation that remains unexplained in any kinetoplastid (111,112). R-loops have been implicated in unscheduled, origin-independent DNA replication at rRNA genes in yeast (38). Such a reaction would not be detected by MFA-seq mapping, which relies on isolation of S phase cells. Thus, we cannot exclude that R-loops can dictate previously undetected origin-independent DNA replication in *T. brucei*, or in other kinetoplastids. Intriguingly, though R-loops or ORC1/CDC6 binding have not been mapped to date in *Leishmania*, very widespread DNA replication initiation events, not focused on the SSRs, have been described throughout each chromosome and appear to localise at intra-CDS regions within PTUs, perhaps indicating a correlation with R-loops (113).

Repeat sequences are well-established loci for the accumulation of R-loops, and several examples of this association are seen in *T. brucei*. First, we detect DRIP-seq enrichment at the parasite’s rRNA loci, as seen in both eukaryotes (81,114) and bacteria (115); as the signal is most pronounced, both in WT cells and in *Tbrh1*−/− mutants, in intra-CDS sequences, it seems likely R-loops form during transcription and not by binding of the mature RNA to DNA. Second, we detect R-loop signal at RHS-rich loci, with a dramatic increase in signal after loss of TbRH1. Here, the genesis of the R-loops here is unclear; for instance, do they arise during RHS transcription, or might they be due to siRNA detected at RHS-associated retrotransposon sequences (106)?. Third, we show that repeat sequences within the mapped centromeres of *T. brucei* are a very pronounced location for R-loop formation. In the three chromosomes where R-loops co-localise with centromeres containing 147 bp repeats, siRNAs may provide the RNA component of the hybrids (106). However, siRNA has not been detected at five chromosomes with centromeres that possess distinct, non-147 bp repeats (106,116), and so the genesis of R-loops at these loci is unclear. Nonetheless, R-loop association with apparently all *T. brucei* centromeres appears to reveal a conserved function for these DNA elements, perhaps related to chromatin modification (117) or cell cycle signalling (118), despite pronounced divergence of the kinetoplastid kinetochore complex that binds centromeres (119,120). Finally, previous work in *T. brucei* (121) and *Leishmania* (122) has revealed the existence of TERRA RNA, with over expression of TbRH1 suppressing the levels of R-loops at *T. brucei* telomere repeats. However, the extent to which (sub)telomeric R-loops might contribute to kinetoplastid biology, such as the expression or stability of telomere-proximal genes, has been little explored.

## Conclusion

The unusual genetic landscape of the *T. brucei* genome, comprising multigene transcription units bounded by a relatively small number of promoters and replication origins, provides insight into the functions of R-loops in genome biology. Repeat sequences are a conserved location of RNA-DNA hybrid formation. In contrast, the novel strategy used by *T. brucei* (and all kinetoplastids) for to gene expression is reflected in potentially novel roles for R-loops in the processing of pre-mRNA transcripts and in transcription initiation at dispersed, unregulated RNA Pol II promoters. Perhaps surprisingly, R-loop mapping provides little evidence that RNA-DNA hybrids contribute to transcription termination or origin-directed DNA replication mapping in *T. brucei*.

## Data Access

Sequences used in the mapping have been deposited in the European Nucleotide Archive (accession number PRJEB21868). DRIPseq analysis will be hosted at TriTryDB (http://tritrypdb.org/tritrypdb/) in an upcoming release.

## Acknowledgements

We thank David Horn and Sebastian Hutchinson, and all lab members, for helpful discussions of emerging data.

## Supplementary data

Available online, comprising 11 figures and one table.

## References

1. Shaw, N.N. and Arya, D.P. (2008) Recognition of the unique structure of DNA:RNA hybrids. Biochimie, 90, 1026–1039.

2. Roberts, R.W. and Crothers, D.M. (1992) Stability and properties of double and triple helices: dramatic effects of RNA or DNA backbone composition. Science, 258, 1463–1466.

3. Dominguez-Sanchez, M.S., Barroso, S., Gomez-Gonzalez, B., Luna, R. and Aguilera, A. (2011) Genome instability and transcription elongation impairment in human cells depleted of THO/TREX. PLoS genetics, 7, e1002386.

4. Huertas, P. and Aguilera, A. (2003) Cotranscriptionally formed DNA:RNA hybrids mediate transcription elongation impairment and transcription-associated recombination. Molecular cell, 12, 711–721.

5. Li, X. and Manley, J.L. (2005) Inactivation of the SR protein splicing factor ASF/SF2 results in genomic instability. Cell, 122, 365–378.

6. Wahba, L., Amon, J.D., Koshland, D. and Vuica-Ross, M. (2011) RNase H and multiple RNA biogenesis factors cooperate to prevent RNA:DNA hybrids from generating genome instability. Molecular cell, 44, 978–988.

7. Stirling, P.C., Chan, Y.A., Minaker, S.W., Aristizabal, M.J., Barrett, I., Sipahimalani, P., Kobor, M.S. and Hieter, P. (2012) R-loop-mediated genome instability in mRNA cleavage and polyadenylation mutants. Genes & development, 26, 163–175.

8. Santos-Pereira, J.M. and Aguilera, A. (2015) R loops: new modulators of genome dynamics and function. Nature reviews. Genetics, 16, 583–597.

9. Costantino, L. and Koshland, D. (2015) The Yin and Yang of R-loop biology. Current opinion in cell biology, 34, 39–45.

10. Skourti-Stathaki, K. and Proudfoot, N.J. (2014) A double-edged sword: R loops as threats to genome integrity and powerful regulators of gene expression. Genes & development, 28, 1384–1396.

11. Koonin, E.V., Makarova, K.S. and Zhang, F. (2017) Diversity, classification and evolution of CRISPR-Cas systems. Current opinion in microbiology, 37, 67–78.

12. Wahba, L., Gore, S.K. and Koshland, D. (2013) The homologous recombination machinery modulates the formation of RNA-DNA hybrids and associated chromosome instability. eLife, 2, e00505.

13. Keskin, H., Shen, Y., Huang, F., Patel, M., Yang, T., Ashley, K., Mazin, A.V. and Storici, F. (2014) Transcript-RNA-templated DNA recombination and repair. Nature, 515, 436–439.

14. Mazina, O.M., Keskin, H., Hanamshet, K., Storici, F. and Mazin, A.V. (2017) Rad52 Inverse Strand Exchange Drives RNA-Templated DNA Double-Strand Break Repair. Molecular cell, 67, 19–29 e13.

15. Zaitsev, E.N. and Kowalczykowski, S.C. (2000) A novel pairing process promoted by Escherichia coli RecA protein: inverse DNA and RNA strand exchange. Genes & development, 14, 740–749.

16. Stirling, P.C. and Hieter, P. (2016) Canonical DNA Repair Pathways Influence R-Loop-Driven Genome Instability. Journal of molecular biology.

17. Sollier, J. and Cimprich, K.A. (2015) Breaking bad: R-loops and genome integrity. Trends in cell biology.

18. Aguilera, A. and Garcia-Muse, T. (2012) R loops: from transcription byproducts to threats to genome stability. Molecular cell, 46, 115–124.

19. Lang, K.S., Hall, A.N., Merrikh, C.N., Ragheb, M., Tabakh, H., Pollock, A.J., Woodward, J.J., Dreifus, J.E. and Merrikh, H. (2017) Replication-Transcription Conflicts Generate R-Loops that Orchestrate Bacterial Stress Survival and Pathogenesis. Cell, 170, 787–799 e718.

20. Hamperl, S., Bocek, M.J., Saldivar, J.C., Swigut, T. and Cimprich, K.A. (2017) Transcription-Replication Conflict Orientation Modulates R-Loop Levels and Activates Distinct DNA Damage Responses. Cell, 170, 774–786 e719.

21. Hamperl, S. and Cimprich, K.A. (2014) The contribution of co-transcriptional RNA:DNA hybrid structures to DNA damage and genome instability. DNA Repair (Amst), 19, 84–94.

22. Cerritelli, S.M. and Crouch, R.J. (2009) Ribonuclease H: the enzymes in eukaryotes. The FEBS journal, 276, 1494–1505.

23. Cerritelli, S.M. and Crouch, R.J. (2016) The Balancing Act of Ribonucleotides in DNA. Trends in biochemical sciences, 41, 434–445.

24. El Hage, A., Webb, S., Kerr, A. and Tollervey, D. (2014) Genome-wide distribution of RNA-DNA hybrids identifies RNase H targets in tRNA genes, retrotransposons and mitochondria. PLoS genetics, 10, e1004716.

25. Cerritelli, S.M., Frolova, E.G., Feng, C., Grinberg, A., Love, P.E. and Crouch, R.J. (2003) Failure to produce mitochondrial DNA results in embryonic lethality in Rnaseh1 null mice. Molecular cell, 11, 807–815.

26. Yang, Z., Hou, Q., Cheng, L., Xu, W., Hong, Y., Li, S. and Sun, Q. (2017) RNase H1 Cooperates with DNA Gyrases to Restrict R-Loops and Maintain Genome Integrity in Arabidopsis Chloroplasts. Plant Cell, 29, 2478–2497.

27. Reijns, M.A., Rabe, B., Rigby, R.E., Mill, P., Astell, K.R., Lettice, L.A., Boyle, S., Leitch, A., Keighren, M., Kilanowski, F. et al. (2012) Enzymatic removal of ribonucleotides from DNA is essential for mammalian genome integrity and development. Cell, 149, 1008–1022.

28. Chan, Y.A., Aristizabal, M.J., Lu, P.Y., Luo, Z., Hamza, A., Kobor, M.S., Stirling, P.C. and Hieter, P. (2014) Genome-wide profiling of yeast DNA:RNA hybrid prone sites with DRIP-chip. PLoS genetics, 10, e1004288.

29. Schwab, R.A., Nieminuszczy, J., Shah, F., Langton, J., Lopez Martinez, D., Liang, C.C., Cohn, M.A., Gibbons, R.J., Deans, A.J. and Niedzwiedz, W. (2015) The Fanconi Anemia Pathway Maintains Genome Stability by Coordinating Replication and Transcription. Molecular cell, 60, 351–361.

30. Garcia-Rubio, M.L., Perez-Calero, C., Barroso, S.I., Tumini, E., Herrera-Moyano, E., Rosado, I.V. and Aguilera, A. (2015) The Fanconi Anemia Pathway Protects Genome Integrity from R-loops. PLoS genetics, 11, e1005674.

31. Bhatia, V., Barroso, S.I., Garcia-Rubio, M.L., Tumini, E., Herrera-Moyano, E. and Aguilera, A. (2014) BRCA2 prevents R-loop accumulation and associates with TREX-2 mRNA export factor PCID2. Nature, 511, 362–365.

32. Tracy, R.B. and Lieber, M.R. (2000) Transcription-dependent R-loop formation at mammalian class switch sequences. The EMBO journal, 19, 1055–1067.

33. Wiedemann, E.M., Peycheva, M. and Pavri, R. (2016) DNA Replication Origins in Immunoglobulin Switch Regions Regulate Class Switch Recombination in an R-Loop-Dependent Manner. Cell reports, 17, 2927–2942.

34. Lilly, J. and Camps, M. (2015) Mechanisms of Theta Plasmid Replication. Microbiology spectrum, 3.

35. Bailey, L.J. and Doherty, A.J. (2017) Mitochondrial DNA replication: a PrimPol perspective. Biochemical Society transactions, 45, 513–529.

36. Noble, E., Spiering, M.M. and Benkovic, S.J. (2015) Coordinated DNA Replication by the Bacteriophage T4 Replisome. Viruses, 7, 3186–3200.

37. Maduike, N.Z., Tehranchi, A.K., Wang, J.D. and Kreuzer, K.N. (2014) Replication of the Escherichia coli chromosome in RNase HI-deficient cells: multiple initiation regions and fork dynamics. Molecular microbiology, 91, 39–56.

38. Stuckey, R., Garcia-Rodriguez, N., Aguilera, A. and Wellinger, R.E. (2015) Role for RNA:DNA hybrids in origin-independent replication priming in a eukaryotic system. Proceedings of the National Academy of Sciences of the United States of America, 112, 5779–5784.

39. Lombrana, R., Almeida, R., Alvarez, A. and Gomez, M. (2015) R-loops and initiation of DNA replication in human cells: a missing link? Frontiers in genetics, 6, 158.

40. Ginno, P.A., Lott, P.L., Christensen, H.C., Korf, I. and Chedin, F. (2012) R-loop formation is a distinctive characteristic of unmethylated human CpG island promoters. Molecular cell, 45, 814–825.

41. Sanz, L.A., Hartono, S.R., Lim, Y.W., Steyaert, S., Rajpurkar, A., Ginno, P.A., Xu, X. and Chedin, F. (2016) Prevalent, Dynamic, and Conserved R-Loop Structures Associate with Specific Epigenomic Signatures in Mammals. Molecular cell, 63, 167–178.

42. Boque-Sastre, R., Soler, M., Oliveira-Mateos, C., Portela, A., Moutinho, C., Sayols, S., Villanueva, A., Esteller, M. and Guil, S. (2015) Head-to-head antisense transcription and R-loop formation promotes transcriptional activation. Proceedings of the National Academy of Sciences of the United States of America, 112, 5785–5790.

43. Sun, Q., Csorba, T., Skourti-Stathaki, K., Proudfoot, N.J. and Dean, C. (2013) R-loop stabilization represses antisense transcription at the Arabidopsis FLC locus. Science, 340, 619–621.

44. Skourti-Stathaki, K., Kamieniarz-Gdula, K. and Proudfoot, N.J. (2014) R-loops induce repressive chromatin marks over mammalian gene terminators. Nature, 516, 436–439.

45. Skourti-Stathaki, K., Proudfoot, N.J. and Gromak, N. (2011) Human senataxin resolves RNA/DNA hybrids formed at transcriptional pause sites to promote Xrn2-dependent termination. Molecular cell, 42, 794–805.

46. Grzechnik, P., Gdula, M.R. and Proudfoot, N.J. (2015) Pcf11 orchestrates transcription termination pathways in yeast. Genes & development, 29, 849–861.

47. Ginno, P.A., Lim, Y.W., Lott, P.L., Korf, I. and Chedin, F. (2013) GC skew at the 5′ and 3′ ends of human genes links R-loop formation to epigenetic regulation and transcription termination. Genome research, 23, 1590–1600.

48. Chedin, F. (2016) Nascent Connections: R-Loops and Chromatin Patterning. Trends in genetics : TIG, 32, 828–838.

49. Zhang, H., Gan, H., Wang, Z., Lee, J.H., Zhou, H., Ordog, T., Wold, M.S., Ljungman, M. and Zhang, Z. (2017) RPA Interacts with HIRA and Regulates H3.3 Deposition at Gene Regulatory Elements in Mammalian Cells. Molecular cell, 65, 272–284.

50. Al-Hadid, Q. and Yang, Y. (2016) R-loop: an emerging regulator of chromatin dynamics. Acta Biochim Biophys Sin (Shanghai), 48, 623–631.

51. Xu, W., Xu, H., Li, K., Fan, Y., Liu, Y., Yang, X. and Sun, Q. (2017) The R-loop is a common chromatin feature of the Arabidopsis genome. Nat Plants.

52. Wahba, L., Costantino, L., Tan, F.J., Zimmer, A. and Koshland, D. (2016) S1-DRIP-seq identifies high expression and polyA tracts as major contributors to R-loop formation. Genes & development, 30, 1327–1338.

53. Rippe, K. and Luke, B. (2015) TERRA and the state of the telomere. Nature structural & molecular biology, 22, 853–858.

54. Kar, A., Willcox, S. and Griffith, J.D. (2016) Transcription of telomeric DNA leads to high levels of homologous recombination and t-loops. Nucleic Acids Res, 44, 9369–9380.

55. Yu, T.Y., Kao, Y.W. and Lin, J.J. (2014) Telomeric transcripts stimulate telomere recombination to suppress senescence in cells lacking telomerase. Proceedings of the National Academy of Sciences of the United States of America, 111, 3377–3382.

56. Clayton, C.E. (2016) Gene expression in Kinetoplastids. Current opinion in microbiology, 32, 46–51.

57. Daniels, J.P., Gull, K. and Wickstead, B. (2010) Cell biology of the trypanosome genome. Microbiol.Mol.Biol.Rev., 74, 552–569.

58. Chikne, V., Gupta, S.K., Doniger, T., k,S.R., Cohen-Chalamish, S., Ben-Asher, H.W., Kolet, L., Yahia, N.H., Unger, R., Ullu, E. et al. (2017) The Canonical Poly (A) Polymerase PAP1 Polyadenylates NonCoding RNAs and Is Essential for snoRNA Biogenesis in Trypanosoma brucei. Journal of molecular biology, 429, 3301–3318.

59. Siegel, T.N., Gunasekera, K., Cross, G.A. and Ochsenreiter, T. (2011) Gene expression in Trypanosoma brucei: lessons from high-throughput RNA sequencing. Trends Parasitol., 27, 434–441.

60. Gunzl, A., Bruderer, T., Laufer, G., Schimanski, B., Tu, L.C., Chung, H.M., Lee, P.T. and Lee, M.G. (2003) RNA polymerase I transcribes procyclin genes and variant surface glycoprotein gene expression sites in Trypanosoma brucei. Eukaryot.Cell, 2, 542–551.

61. Kolev, N.G., Franklin, J.B., Carmi, S., Shi, H., Michaeli, S. and Tschudi, C. (2010) The transcriptome of the human pathogen Trypanosoma brucei at single-nucleotide resolution. PLoS pathogens, 6, e1001090.

62. Siegel, T.N., Hekstra, D.R., Kemp, L.E., Figueiredo, L.M., Lowell, J.E., Fenyo, D., Wang, X., Dewell, S. and Cross, G.A. (2009) Four histone variants mark the boundaries of polycistronic transcription units in Trypanosoma brucei. Genes Dev., 23, 1063–1076.

63. Wedel, C., Forstner, K.U., Derr, R. and Siegel, T.N. (2017) GT-rich promoters can drive RNA pol II transcription and deposition of H2A.Z in African trypanosomes. The EMBO journal.

64. Fadda, A., Ryten, M., Droll, D., Rojas, F., Farber, V., Haanstra, J.R., Merce, C., Bakker, B.M., Matthews, K. and Clayton, C. (2014) Transcriptome-wide analysis of trypanosome mRNA decay reveals complex degradation kinetics and suggests a role for co-transcriptional degradation in determining mRNA levels. Molecular microbiology, 94, 307–326.

65. Reynolds, D., Hofmeister, B.T., Cliffe, L., Alabady, M., Siegel, T.N., Schmitz, R.J. and Sabatini, R.(2016) Histone H3 Variant Regulates RNA Polymerase II Transcription Termination and Dual Strand Transcription of siRNA Loci in Trypanosoma brucei. PLoS genetics, 12, e1005758.

66. Schulz, D., Zaringhalam, M., Papavasiliou, F.N. and Kim, H.S. (2016) Base J and H3.V Regulate Transcriptional Termination in Trypanosoma brucei. PLoS genetics, 12, e1005762.

67. Tiengwe, C., Marcello, L., Farr, H., Dickens, N., Kelly, S., Swiderski, M., Vaughan, D., Gull, K., Barry, J.D., Bell, S.D. et al. (2012) Genome-wide analysis reveals extensive functional interaction between DNA replication initiation and transcription in the genome of Trypanosoma brucei. Cell reports, 2, 185–197.

68. Devlin, R., Marques, C.A., Paape, D., Prorocic, M., Zurita-Leal, A.C., Campbell, S.J., Lapsley, C., Dickens, N. and McCulloch, R. (2016) Mapping replication dynamics in Trypanosoma brucei reveals a link with telomere transcription and antigenic variation. eLife, 5.

69. Marques, C.A., Tiengwe, C., Lemgruber, L., Damasceno, J.D., Scott, A., Paape, D., Marcello, L. and McCulloch, R. (2016) Diverged composition and regulation of the Trypanosoma brucei origin recognition complex that mediates DNA replication initiation. Nucleic Acids Res, 44, 4763–4784.

70. Tiengwe, C., Marques, C.A. and McCulloch, R. (2014) Nuclear DNA replication initiation in kinetoplastid parasites: new insights into an ancient process. Trends in parasitology, 30, 27–36.

71. Boguslawski, S.J., Smith, D.E., Michalak, M.A., Mickelson, K.E., Yehle, C.O., Patterson, W.L. and Carrico, R.J. (1986) Characterization of monoclonal antibody to DNA.RNA and its application to immunodetection of hybrids. J Immunol Methods, 89, 123–130.

72. Garcia-Rubio, M., Barroso, S.I. and Aguilera, A. (2018) Detection of DNA-RNA Hybrids In Vivo. Methods in molecular biology, 1672, 347–361.

73. Hutchinson, S., Glover, L. and Horn, D. (2016) High-resolution analysis of multi-copy variant surface glycoprotein gene expression sites in African trypanosomes. BMC Genomics, 17, 806.

74. Langmead, B. and Salzberg, S.L. (2012) Fast gapped-read alignment with Bowtie 2. Nature methods, 9, 357–359.

75. Li, H., Handsaker, B., Wysoker, A., Fennell, T., Ruan, J., Homer, N., Marth, G., Abecasis, G. and Durbin, R. (2009) The Sequence Alignment/Map format and SAMtools. Bioinformatics., 25, 2078–2079.

76. Robinson, J.T., Thorvaldsdottir, H., Winckler, W., Guttman, M., Lander, E.S., Getz, G. and Mesirov, J.P. (2011) Integrative genomics viewer. Nat Biotechnol, 29, 24–26.

77. Bailey, T.L., Johnson, J., Grant, C.E. and Noble, W.S. (2015) The MEME Suite. Nucleic Acids Res, 43, W39–49.

78. Ramirez, F., Dundar, F., Diehl, S., Gruning, B.A. and Manke, T. (2014) deepTools: a flexible platform for exploring deep-sequencing data. Nucleic Acids Res, 42, W187–191.

79. Siegel, T.N., Hekstra, D.R., Wang, X., Dewell, S. and Cross, G.A. (2010) Genome-wide analysis of mRNA abundance in two life-cycle stages of Trypanosoma brucei and identification of splicing and polyadenylation sites. Nucleic Acids Res, 38, 4946–4957.

80. Halasz, L., Karanyi, Z., Boros-Olah, B., Kuik-Rozsa, T., Sipos, E., Nagy, E., Mosolygo, L.A., Mazlo, A., Rajnavolgyi, E., Halmos, G. et al. (2017) RNA-DNA hybrid (R-loop) immunoprecipitation mapping: an analytical workflow to evaluate inherent biases. Genome research, 27, 1063–1073.

81. El Hage, A., French, S.L., Beyer, A.L. and Tollervey, D. (2010) Loss of Topoisomerase I leads to R-loop-mediated transcriptional blocks during ribosomal RNA synthesis. Genes & development, 24, 1546–1558.

82. Salvi, J.S., Chan, J.N., Szafranski, K., Liu, T.T., Wu, J.D., Olsen, J.B., Khanam, N., Poon, B.P., Emili, A. and Mekhail, K. (2014) Roles for Pbp1 and caloric restriction in genome and lifespan maintenance via suppression of RNA-DNA hybrids. Dev Cell, 30, 177–191.

83. Ohle, C., Tesorero, R., Schermann, G., Dobrev, N., Sinning, I. and Fischer, T. (2016) Transient RNA-DNA Hybrids Are Required for Efficient Double-Strand Break Repair. Cell, 167, 1001–1013 e1007.

84. Yang, Y., La, H., Tang, K., Miki, D., Yang, L., Wang, B., Duan, C.G., Nie, W., Wang, X., Wang, S. et al. (2017) SAC3B, a central component of the mRNA export complex TREX-2, is required for prevention of epigenetic gene silencing in Arabidopsis. Nucleic Acids Res, 45, 181–197.

85. Berriman, M., Ghedin, E., Hertz-Fowler, C., Blandin, G., Renauld, H., Bartholomeu, D.C., Lennard, N.J., Caler, E., Hamlin, N.E., Haas, B. et al. (2005) The genome of the African trypanosome Trypanosoma brucei. Science, 309, 416–422.

86. Jenjaroenpun, P., Wongsurawat, T., Yenamandra, S.P. and Kuznetsov, V.A. (2015) QmRLFS-finder: a model, web server and stand-alone tool for prediction and analysis of R-loop forming sequences. Nucleic Acids Res, 43, W527–534.

87. Benson, G. (1999) Tandem repeats finder: a program to analyze DNA sequences. Nucleic Acids Res, 27, 573–580.

88. Echeverry, M.C., Bot, C., Obado, S.O., Taylor, M.C. and Kelly, J.M. (2012) Centromere-associated repeat arrays on Trypanosoma brucei chromosomes are much more extensive than predicted. BMC Genomics, 13, 29.

89. Savage, A.F., Kolev, N.G., Franklin, J.B., Vigneron, A., Aksoy, S. and Tschudi, C. (2016) Transcriptome Profiling of Trypanosoma brucei Development in the Tsetse Fly Vector Glossina morsitans. PLoS One, 11, e0168877.

90. Dunbar, D.A., Chen, A.A., Wormsley, S. and Baserga, S.J. (2000) The genes for small nucleolar RNAs in Trypanosoma brucei are organized in clusters and are transcribed as a polycistronic RNA. Nucleic Acids Res, 28, 2855–2861.

91. Moraes Barros, R.R., Marini, M.M., Antonio, C.R., Cortez, D.R., Miyake, A.M., Lima, F.M., Ruiz, J.C., Bartholomeu, D.C., Chiurillo, M.A., Ramirez, J.L. et al. (2012) Anatomy and evolution of telomeric and subtelomeric regions in the human protozoan parasite Trypanosoma cruzi. BMC Genomics, 13, 229.

92. Bringaud, F., Biteau, N., Melville, S.E., Hez, S., El Sayed, N.M., Leech, V., Berriman, M., Hall, N., Donelson, J.E. and Baltz, T. (2002) A new, expressed multigene family containing a hot spot for insertion of retroelements is associated with polymorphic subtelomeric regions of Trypanosoma brucei. Eukaryot.Cell, 1, 137–151.

93. Siegel, T.N., Hekstra, D.R., Wang, X., Dewell, S. and Cross, G.A. (2010) Genome-wide analysis of mRNA abundance in two life-cycle stages of Trypanosoma brucei and identification of splicing and polyadenylation sites. Nucleic Acids Res., 38, 4946–4957.

94. Tiengwe, C., Marcello, L., Farr, H., Gadelha, C., Burchmore, R., Barry, J.D., Bell, S.D. and McCulloch, R. (2012) Identification of ORC1/CDC6-Interacting Factors in Trypanosoma brucei Reveals Critical Features of Origin Recognition Complex Architecture. PLoS One, 7, e32674.

95. Michaeli, S. (2011) Trans-splicing in trypanosomes: machinery and its impact on the parasite transcriptome. Future Microbiol, 6, 459–474.

96. Clayton, C. and Michaeli, S. (2011) 3′ processing in protists. Wiley Interdiscip Rev RNA, 2, 247–255.

97. Koch, H., Raabe, M., Urlaub, H., Bindereif, A. and Preusser, C. (2016) The polyadenylation complex of Trypanosoma brucei: Characterization of the functional poly(A) polymerase. RNA biology, 13, 221–231.

98. Matthews, K.R., Tschudi, C. and Ullu, E. (1994) A common pyrimidine-rich motif governs transsplicing and polyadenylation of tubulin polycistronic pre-mRNA in trypanosomes. Genes & development, 8, 491–501.

99. Ullu, E., Matthews, K.R. and Tschudi, C. (1993) Temporal order of RNA-processing reactions in trypanosomes: rapid trans splicing precedes polyadenylation of newly synthesized tubulin transcripts. Mol.Cell Biol., 13, 720–725.

100. Bonnet, A., Grosso, A.R., Elkaoutari, A., Coleno, E., Presle, A., Sridhara, S.C., Janbon, G., Geli, V., de Almeida, S.F. and Palancade, B. (2017) Introns Protect Eukaryotic Genomes from Transcription-Associated Genetic Instability. Molecular cell, 67, 608–621 e606.

101. Dumelie, J.G. and Jaffrey, S.R. (2017) Defining the location of promoter-associated R-loops at near-nucleotide resolution using bisDRIP-seq. eLife, 6.

102. Chen, L., Chen, J.Y., Zhang, X., Gu, Y., Xiao, R., Shao, C., Tang, P., Qian, H., Luo, D., Li, H. et al. (2017) R-ChIP Using Inactive RNase H Reveals Dynamic Coupling of R-loops with Transcriptional Pausing at Gene Promoters. Molecular cell, 68, 745–757 e745.

103. Schwartz, S. and Ast, G. (2010) Chromatin density and splicing destiny: on the cross-talk between chromatin structure and splicing. The EMBO journal, 29, 1629–1636.

104. Lenhard, B., Sandelin, A. and Carninci, P. (2012) Metazoan promoters: emerging characteristics and insights into transcriptional regulation. Nature reviews. Genetics, 13, 233–245.

105. Li, H., Hou, J., Bai, L., Hu, C., Tong, P., Kang, Y., Zhao, X. and Shao, Z. (2015) Genome-wide analysis of core promoter structures in Schizosaccharomyces pombe with DeepCAGE. RNA biology, 12, 525–537.

106. Tschudi, C., Shi, H., Franklin, J.B. and Ullu, E. (2012) Small interfering RNA-producing loci in the ancient parasitic eukaryote Trypanosoma brucei. BMC Genomics, 13, 427.

107. Powell, W.T., Coulson, R.L., Gonzales, M.L., Crary, F.K., Wong, S.S., Adams, S., Ach, R.A., Tsang, P., Yamada, N.A., Yasui, D.H. et al. (2013) R-loop formation at Snord116 mediates topotecan inhibition of Ube3a-antisense and allele-specific chromatin decondensation. Proceedings of the National Academy of Sciences of the United States of America, 110, 13938–13943.

108. Mischo, H.E., Gomez-Gonzalez, B., Grzechnik, P., Rondon, A.G., Wei, W., Steinmetz, L., Aguilera, A. and Proudfoot, N.J. (2011) Yeast Sen1 helicase protects the genome from transcription-associated instability. Molecular cell, 41, 21–32.

109. Reynolds, D., Cliffe, L., Forstner, K.U., Hon, C.C., Siegel, T.N. and Sabatini, R. (2014) Regulation of transcription termination by glucosylated hydroxymethyluracil, base J, in Leishmania major and Trypanosoma brucei. Nucleic Acids Res, 42, 9717–9729.

110. van Luenen, H.G., Farris, C., Jan, S., Genest, P.A., Tripathi, P., Velds, A., Kerkhoven, R.M., Nieuwland, M., Haydock, A., Ramasamy, G. et al. (2012) Glucosylated hydroxymethyluracil, DNA base J, prevents transcriptional readthrough in Leishmania. Cell, 150, 909–921.

111. Marques, C.A.a.M., R. (2018) Conservation and Variation in Strategies for DNA Replication of Kinetoplastid Nuclear Genomes. Current genomics, 19, 98–109.

112. da Silva, M.S., Pavani, R.S., Damasceno, J.D., Marques, C.A., McCulloch, R., Tosi, L.R.O. and Elias, M.C. (2017) Nuclear DNA Replication in Trypanosomatids: There Are No Easy Methods for Solving Difficult Problems. Trends in parasitology, 33, 858–874.

113. Lombrana, R., Alvarez, A., Fernandez-Justel, J.M., Almeida, R., Poza-Carrion, C., Gomes, F., Calzada, A., Requena, J.M. and Gomez, M. (2016) Transcriptionally Driven DNA Replication Program of the Human Parasite Leishmania major. Cell reports, 16, 1774–1786.

114. Arana, M.E., Kerns, R.T., Wharey, L., Gerrish, K.E., Bushel, P.R. and Kunkel, T.A. (2012) Transcriptional responses to loss of RNase H2 in Saccharomyces cerevisiae. DNA Repair (Amst), 11, 933–941.

115. Hraiky, C., Raymond, M.A. and Drolet, M. (2000) RNase H overproduction corrects a defect at the level of transcription elongation during rRNA synthesis in the absence of DNA topoisomerase I in Escherichia coli. The Journal of biological chemistry, 275, 11257–11263.

116. Obado, S.O., Bot, C., Nilsson, D., Andersson, B. and Kelly, J.M. (2007) Repetitive DNA is associated with centromeric domains in Trypanosoma brucei but not Trypanosoma cruzi. Genome Biol., 8, R37.

117. Castellano-Pozo, M., Santos-Pereira, J.M., Rondon, A.G., Barroso, S., Andujar, E., Perez-Alegre, M., Garcia-Muse, T. and Aguilera, A. (2013) R loops are linked to histone H3 S10 phosphorylation and chromatin condensation. Molecular cell, 52, 583–590.

118. Kabeche, L., Nguyen, H.D., Buisson, R. and Zou, L. (2018) A mitosis-specific and R loop-driven ATR pathway promotes faithful chromosome segregation. Science, 359, 108–114.

119. Akiyoshi, B. and Gull, K. (2014) Discovery of unconventional kinetochores in kinetoplastids. Cell, 156, 1247–1258.

120. D’Archivio, S. and Wickstead, B. (2016) Trypanosome outer kinetochore proteins suggest conservation of chromosome segregation machinery across eukaryotes. The Journal of cell biology.

121. Nanavaty, V., Sandhu, R., Jehi, S.E., Pandya, U.M. and Li, B. (2017) Trypanosoma brucei RAP1 maintains telomere and subtelomere integrity by suppressing TERRA and telomeric RNA:DNA hybrids. Nucleic Acids Res.

122. Damasceno, J.D., Silva, G., Tschudi, C. and Tosi, L.R. (2017) Evidence for regulated expression of Telomeric Repeat-containing RNAs (TERRA) in parasitic trypanosomatids. Mem Inst Oswaldo Cruz, 112, 572–576.

